# H3K27me3 mediated KRT14 upregulation promotes TNBC peritoneal metastasis

**DOI:** 10.1101/2021.02.02.429349

**Authors:** Ayushi Verma, Akhilesh Singh, Mushtaq Ahmad Nengroo, Krishan Kumar Saini, Abhipsa Sinha, Anup Kumar Singh, Dipak Datta

## Abstract

Triple Negative Breast Cancer (TNBC) is known to have poor prognosis and adverse clinical outcome among all breast cancer subtypes due to the absence of available targeted therapy for it. Emerging literature indicates that epigenetic reprogramming is now appreciated as a driving force for TNBC pathophysiology. High expression of epigenetic modulator EZH2 (Enhancer of zeste homolog 2) has been shown to correlate with TNBC poor prognosis but the contribution of EZH2 catalytic (H3K27me3) versus non-catalytic EZH2 (NC-EZH2) function in TNBC growth and progression remains elusive. In the process of dissecting the impact of H3K27me3 versus NC-EZH2 function in TNBC pathogenesis, we reveal that selective hyperactivation of H3K27me3 over NC-EZH2 not only promotes TNBC metastasis but also alters the metastatic landscape of TNBC. Using extensive *in- vivo* live animal imaging, we present conclusive evidence that peritoneal metastasis, particularly splenic metastasis of TNBC is governed by H3K27me3. Transcriptome analyses of hyperactive H3K27me3 cells lead us to discover Cytokeratin-14 (KRT14) as a new target of H3K27me3. Unlike classical H3K27me3 mediated suppression of gene expression, here; we observe that H3K27me3 enhances KRT14 transcription by attenuating the binding of transcriptional repressor Sp1 to its promoter. Further, loss of KRT14 significantly reduces TNBC migration, invasion and splenic metastasis. Finally, genetic ablation of EZH2 or pharmacological inhibition of EZH2 catalytic function by FDA approved drug tazemetostat (EPZ6438) robustly inhibits TNBC peritoneal metastasis. Altogether, our preclinical findings posit a rational insight for the clinical development of H3K27me3 inhibitor like tazemetostat as a targeted therapy against TNBC.

## Introduction

According to global cancer statistics, breast cancer is the most common malignancy worldwide in women, having 2.09 million new cases diagnosed in 2018 with 0.6 million deaths (Bray et al., 2018). Among different breast cancer subtypes, basal-like breast cancer often clinically defined as Triple Negative Breast Cancer (TNBC) which lacks of immunohistochemistry (IHC) expression of Estrogen receptor (ER), progesterone receptor (PR) and human epidermal growth factor receptor 2 (her2/neu) according to College of American Pathologist (CAP)/ASCO guidelines (Wolff et al., 2014). It involves young patient age and is associated with high grade, poor prognosis (Cheang et al., 2008). TNBC poses serious threat to clinicians due to the enormous heterogeneity of the disease and the absence of well-established molecular targets (Lehmann et al., 2011). Indeed, the 90% mortality of breast cancer including TNBC is associated with metastasis (Chaffer and Weinberg, 2011; Dent et al., 2007; Gupta and Massagué, 2006; Kennecke et al., 2010). Underpinning the molecular cues for TNBC metastasis and its therapeutic intervention are in the hotspots of current cancer research.

The concept of epigenetic reprogramming is being admired as a driving force for the distant organ metastasis. Importantly, breast cancer molecular subtypes have been identified with unique chromatin architecture with diverse methylation patterns (Holm et al., 2012; Karsli-Ceppioglu et al., 2017). However, how subtype-specific distinct transcriptional nexus are exclusively regulated by epigenetic mechanisms still remains elusive. Epigenetic modulator Enhancer of Zeste homolog 2 (EZH2), a catalytic subunit of Polycomb Repressive Complex 2 (PRC2), promotes target genes suppression by tri-methylation of lysine 27 of histone H3 (H3K27me3) (Kleer et al., 2003). Recently, we have shown the classical gene silencing function of EZH2 where death receptors are epigenetically suppressed in cancer stem cells (Singh et al., 2020). Notably, aberrant EZH2 expression has been associated with diverse cancers in respect of both oncogenic and tumor suppression functions (Kim and Roberts, 2016). Besides its canonical function as a transcriptional repressor, EZH2 protein has recently been shown to perform H3K27me3 independent functions (Kim et al., 2018). Recently, Mittal’s group has performed the bioinformatics analysis of epigenetic-associated genes in breast cancer patients, which characterized EZH2 overexpression as a predominant carrier in TNBC metastasis with poor overall survival (Yomtoubian et al., 2020).

Expression of basal cytokeratin like CK5/6, CK14 and CK17 are hallmark features of TNBC (Lehmann et al., 2011). Interestingly, cytokeratins serve more than mere epithelial cell marker and several recent studies have provided evidence for active keratin involvement in cancer cell invasion and metastasis (Karantza, 2011). Kevin et.al in their elegant studies showed that polyclonal breast cancer metastases arise from collective dissemination of keratin 14-expressing tumor cell clusters (Cheung et al., 2016). Regulation of cytokeratin expression and their functional consequences particularly in the context to basal like breast cancer metastasis is not clear yet.

In our study, we have dissected the differential role of EZH2 catalytic (H3K27me3) versus non-catalytic EZH2 (NC-EZH2) function in the context of TNBC progression. Interestingly, we identified that hyperactivated H3K27me3 but not its NC-EZH2 protein overexpression resulted in robust increase of TNBC peritoneal metastasis, though primary tumor growth remained unchanged. Transcriptome analysis of H3K27me3 hyperactivated TNBC cells led us to discover that KRT14 is a new target of H3K27me3. Quite intriguingly, instead of H3K27me3 mediated classical transcriptional repression, here, we found H3K27me3 promotes *KRT14* gene expression by altering the recruitment pattern of its transcription factor Sp1. Our sequential *in-vivo* imaging experiments clearly demonstrate that KRT14 is a critical regulator of TNBC splenic metastasis and loss of KRT14 even in H3K27me3 hyper-activated background severely compromised TNBC peritoneal metastasis. Notably, EZH2 functional loss either by its knock down or the treatment of H3K27me3 selective inhibitor EPZ6438 resulted in significant reduction of TNBC migration, invasion *in-vitro* and peritoneal metastasis *in- vivo*. Finally, we discovered clinical relevance of our finding in TCGA database where ER, PR and HER2 negative basal-like breast cancer or TNBC selectively display positive correlation between EZH2 and KRT14 expression. In summary, our data indicate that peritoneal metastasis of TNBC may be driven by the trimethylation function of EZH2, and targeting the methyltransferase activity of EZH2 could be a wonderful strategy to prevent TNBC metastasis in the clinic.

## 2. Materials and Methods

### Reagents and antibodies

Dimethyl Sulfoxide (DMSO), Bovine Serum Albumin (BSA), anti-β-Actin (cat# A3854) antibody, Crystal violet dye, doxycycline and Polybrene were purchased from Sigma-Aldrich. EPZ6438 was purchased from Apex biosciences. XenoLight D-Luciferin potassium salt (P/N 122799) obtained from Perkin Elmer. ProLong^™^ Gold Antifade Mountant was purchased from Invitrogen. Isoflurane (FORANE) was bought from Baxter U.S. Health Care. Matrigel Invasion Chamber (24 well plate 0.8 microns, Lot-8351001) was purchased from Corning. Antibodies for EZH2 (cat#5246 S), H3k27me3 (cat# 9733 S), p-PolII-S5 (cat# 13523 S), Sp1 (# 9389S) and Magnetic ChIP kit were purchased from cell signaling technology (CST). The antibody for GAPDH (#25778) was purchased from Santa Cruz Biotechnology. Antibody for KRT14 procured from abcam (cat# ab7800). PVDF membrane and stripping buffer were obtained from Millipore Inc. BCA protein estimation kit, RIPA cell lysis buffer, blocking buffer, Super Signal West Pico and Femto chemiluminescent substrate, Lipofectamine-3000, Puromycin, Alexafluor 488/594 conjugated secondary antibodies, FBS, RPMI-1640 media, Anti-Anti, DYNAmo Color Flash SYBR Green qPRC kit (cat# F-416L) were purchased from Thermo Fisher Scientific. Primers for real time PCR and ChIP assay were purchased from IDT Inc (Details of primer are listed in the table-1). Iscript AdV cDNA kit for RT-PCR (cat# 1725038) was purchased from *BIO-RAD.* RNeasy Mini Kit (cat#74104) was bought from Qiagen. Ex prep ^™^ plasmid SV mini (cat# 101-150) procured from Gene All. The scrather was purchased from SPL life sciences. The collagenase Type I (cat# 1700-017) was purchased from Gibco. All chemicals and antibodies were obtained from Sigma or Thermo scientific unless specified otherwise.

### Procurement and culture of cell lines

Various basal like or basaloid TNBC human (HCC1806 and MDAMB468) and mouse (4T-1) cell lines were obtained from American Type Culture Collection (ATCC), USA. Mycoplasma free early passage cells were revived from liquid nitrogen vapor stocks and inspected microscopically for stable phenotype before use. Human cell lines used in the study are authenticated by STR profiling. All experiments were performed within early passages (<10) of individual cell lines. Cells were cultured as monolayers in recommended media supplemented with 10% FBS, 1-X anti-anti (containing 100 μg/ml streptomycin, 100 unit/ml penicillin and 0.25 μg/ml amphotericin B) and maintained in 5% CO_2_ and humidified environmental 37°C.

### Generation of stable cell lines

MSCV (cat# 24828), MSCV EZH2 ΔSET-Hygro (cat# 49403) EZH2-Y641-F (cat# 80077) and pTRIPZ M)-YFP-EZH2 (cat# 82511) were procured from Addgene USA. Control (MSCV), EZH2 ΔSET-Hygro and EZH2-Y641-F cell lines were generated by utilizing retroviral mediated transduction system followed by puromycin selection. The HEK293-T cell line was used for the generation of retroviral particles following standard protocol. The HEK293-T cells were plated in the 6-well plate at 80% confluency. Polybrene (8μg/ml) was added to the viral soup during the transduction matured viral particles into the target cells. MSCV control cell were subjected to puromycin and EZH2 Y641-F and EZH2 ΔSET cells were subjected to hygromycin selection, and the overexpression of stable EZH2, ΔSET and EZH2-Y641-F were confirmed by western blot. EZH2 mouse, KRT14 mouse and Human ShRNA sequence were cloned in the 3rd generation transfer plasmid pLKO.1 TRC cloning vector (Addgene cat # 10878) between unique AgeI and EcoRI restriction sites downstream of the U6 promoter. HEK-293T cell line was used for the generation of lentiviral particles and media containing the viral particles was supplemented with Polybrene (8μg/ml) for the transduction purpose. Cells were subjected to puromycin selection after 48 hours of transduction, and the knockdown for EZH2 and KRT14 were confirmed by western blot. ShRNA sequences were listed in the Table-S5.

### Western blotting

Cells were subjected for lysis in RIPA buffer containing phosphatase and protease inhibitor cocktail and incubated at −20°C for 48 hour and subsequently the samples were thawed at RT and centrifuged at 5000g for 15 min at 4° C, supernatant was collected and the pallet was discarded. The Protein concentrations were estimated by utilizing the BCA kit. Equal amounts of protein were resolved by SDS-PAGE and transferred to a PVDF membrane. Membranes were blocked with 5% nonfat dry milk or 5% BSA followed by incubation with appropriate dilutions (1:1000) of primary antibodies overnight at 4°C and subsequently incubated with a 1:5000 dilution of horseradish peroxidase-conjugated secondary antibodies for 1 hour at room temperature. Immunoreactivity was detected by enhanced chemiluminescence solution (ImmobilonTM western, Millipore, USA) and scanned by the gel documentation system (Bio-Rad chemidoc XRS plus).

### RNA Seq analysis

RNA-Seq libraries were created using TruSeq RNA-Seq Library Prep Kit-v2. The experiment was paired-end with 150nt read length, the libraries were sequenced to mean >100X coverage on the illumina sequencing platform. The quality check of raw reads obtained from the next generation sequencing platform was checked using FastQC (v0.11.5) software. Followed by the removal of the adapter (Illumina TruSeq Small RNA 3’Adapter-AGATCGGAAGAGCACACGTCT) using FastX-ToolKit (v0.0.13). Mapping to the reference genome was done using Tophat software (v2.1.1) followed by the abundance estimation using Cufflinks software (v2.2.1) for each sample. All samples transcripts were assembled using the Cuff merge (v2.2.1) and final transcriptome assembly was received as an output. For differential expression estimation Cuffdiff (v2.2.1) was used. The differential expression results obtained from differential expression estimation were visualized using the R language (v3.6.1) DESeq package (v1.36.0) and expression plots were placed.

### Real-time PCR

Total RNA was isolated from cultured cells using the standard procedure of the RNeasy Mini Kit (Qiagen, cat # 74104). The concentration and purity of the RNA samples was determined using nanodrop. Total RNA (7 μg) of each sample was reverse-transcribed (RT) with iscript ADV cDNA synthesis kit. The final cDNA was diluted with nuclease-free water (1:3), 1μl of this having a concentration of 80ng/μl was used for each reaction in real-time PCR. Real-time PCR was carried out using an ABI Step One plus Real-Time PCR System (Applied Biosystems). Reactions for each sample were performed in triplicate. 18s amplification was used as the housekeeping gene. A gene expression score was calculated by taking two raised to the difference in Ct between the housekeeping gene and the gene of interest (2 ΔCt). For amplification of *ATF-3, ID2, ID3, FZD5, KRT8, KRT16, KRT14, FST, DOX7, TLE6, DEPTOR, PTRPUC* and *DLC-1,* we performed SYBR Green based RT-PCR following manufacturer’s instructions. Primers were listed in the Table-S3

### Chromatin Immunoprecipitation (ChIP) Assay

ChIP assay was conducted by using the ChIP assay kit (Cell Signaling Technology) following the manufacturer’s protocol. In brief, cells at 80% confluence were fixed with formaldehyde (1% final concentration directly to the culture media) for 10 minutes. Cells were then centrifuged, followed by lysis in 200μl of membrane extraction buffer containing protease inhibitor cocktail. The cell lysates were digested with MNase for 30 minutes at 37°C to get chromatin fragments followed by sonication (with 20 second on/20 second off 3 Sonication cycles at 50% amplitude) to generate 100-500 bp long DNA fragments. After centrifugation, clear supernatant was diluted (100:400) in 1X ChIP buffer with protease inhibitor cocktail followed by keeping 5% of input control apart and incubated with primary antibody or respective normal IgG antibody overnight at 4°C on a rotor. The next day, IP reactions were incubated for 2 hours in ChIP-Grade Protein G Magnetic Beads, followed by precipitation of beads and sequential washing with a low and high salt solution. Then elution of chromatin from Antibody/Protein G Magnetic beads and reversal of cross-linking was carried out by heat. DNA was purified by using spin columns, and SYBR Green based real-time PCR was conducted. Primer sequences used for the ChIP experiment for different genes are enlisted in Table-S4.

### Confocal microscopy

Control and treated cells were fixed with ice-cold pure methanol for 10 min at −20° C followed by blocking with 2% BSA for 1 hour at RT. After overnight primary antibodies (anti-H3K27me3 and anti-KRT14) incubation, cells were washed twice with PBS and incubated with fluorescent-conjugated secondary antibodies at RT for 1 hour, followed by DAPI staining for 5 min at RT. After washing, cells were mounted with anti-fade mounting medium on glass slides and viewed under an inverted confocal laser scanning microscope (Zeiss Meta 510 LSM; Carl Zeiss, Jena, Germany). Plan Apochromat63X/1.4NA Oil DIC objective lens was used for imaging and data collection. Appropriate excitation lines, excitation and emission filters were used for imaging.

### Wound healing assay

For wound healing assay, 500,000 cells were seeded in to 6 well plates and incubated for overnight to form a confluent monolayer. The scratch has been made by scrather to generate a straight line scratch in the cell monolayer. The cells were washed with PBS and cultured with fresh complete RPMI media at 37°C for different time intervals. The cells were incubated at 37°C for different time intervals with or without treatments. Five (5) reference points were randomly selected from single well at different time intervals and percentage of wounds healed area was measured by Image J software. Three independent replicate experiments were conducted for single data representation.

### Invasion Assay

For each invasion assay, cells were re-suspended in 500μl of serum free RPMI and was added to the inside of Matrigel inserts (Coring Bio Coat) and DMEM media with the 10 % serum was added outsides of inserts. The Matrigel invasion chambers incubated for 24 hours at 37°C. The non-invading cells were then removed by scrubbing with cotton tipped swab. Invasion chambers were fixed with 100 % methanol for 5 minutes. The invasion chambers were stained with crystal violet for 1 hour and then washed with PBS twice. 5 reference points were randomly selected for each invasion camber. The number of invaded cells were analysed by Image J software. Three independent replicate experiments were conducted for single data representation.

### *In -vivo* tumor growth and metastasis studies

All animal studies were conducted by following standard principles and procedures approved by the Institutional Animal Ethics Committee (IAEC) of CSIR-Central Drug Research Institute. For orthotopic inoculation, different tagged (GFP, Td Tomato, Luc) 4T-1 cells (1×10^6^) in 100 μl were injected in to the mammary fat pad of 4-6 week old nude Crl: CD1-Foxn1^nu^ female mice, whereas, same number of cells were inoculated in dorsal right flank for subcutaneous mice model. After 1 week of post inoculation, EPZ6438 (250mg/kg dose) or vehicle (0.5% NaCMC + 0.1% Tween-80 in water) was administered per day by oral gavage for 24 days. Throughout the study, tumors were measured with an electronic digital caliper at regular interval and the tumor volume was calculated using standard formula *V = ⊓ / 6 × a^2^ × b* (*‘a’* is the short and *‘b’* is the long tumor axis). At the end of experiment, mice were sacrificed, and tumors were dissected for further studies. Live animal bioluminescent imaging (IVIS spectrum, Perkin Elmer) performed once per week of post inoculation. For *in- vivo* imaging studies, 150 mg/Kg D-Luciferin (10mg/ml in PBS) was injected intraperitoneal in the tumor bearing mice. Subsequently, mice were anesthetized by Isoflurane. Images were captured with dorsal and ventral positions using Perkin Elmer IVIS system coupled with bioluminescence image acquisition and analysis software. Regions of interest (ROI) from displayed images were identified on the tumor and metastatic sites and quantified as photons per second (p/s) using Living Image software. Spectral unmixing was used to detect Td–tomato and GFP signals in the same mice, and finally data were acquired at 570 nm (Td) and 465 nm (GFP).

### Isolation, culture and analysis of metastatic cells

Mice were sacrificied and primary tumors and other organs were harvested under sterile condition. Single cell suspension was prepared following standard protocol. Briefly, chopped tissues were incubated in HBBS solution containing 1mg/ml of Collagenase on a rocker for 2 hr at 37 °C. Suspension was then centrifuged for 5 min at 500x g, washed and cell pellet was re-suspended in normal growth medium (RPMI, 10% FBS, 1X anti-anti) onto a T-25 flask. After 24 hours, fresh media containing 60 μM 6-Thioguanine was added and cultured for 3 days to keep only 6-Tg resistant 4T1 cells. Isolated cells from primary tumors and metastatic organs were used for further analysis.

### Analysis of TCGA Breast cancer dataset

Illumina HiSeq mRNA data of patients with Breast cancer (Yau 2010) was downloaded from TCGA dataset available at the UCSC Xena (https://xena.ucsc.edu/) for *ESR1, PGR, ERBB2 EZH2* and *KRT14.* The breast cancer dataset was segregated into a PAM50 subtype for luminal A, Luminal B, HER2^+^, basal and normal like subtypes. The heatmap was generated between EZH2 Log2FC values and KRT14 Log2FC values between different breast cancer subtypes.

### Statistics

Most of the *in-vitro* experiments are representative of at least three independent experiments. Two Way ANOVA, One Way ANOVA, Student’s t-test and two-tailed distributions were used to calculate the statistical significance of *in-vitro* and *in-vivo* experiments. The Kaplan Meier survival curve significance was analysed by log rank Test. These analyses were done with Graph Pad Prism software. Results were considered statistically significant when p-values ≤ 0.05 between groups.

## Results

### EZH2 functional activation but not its protein overexpression promotes TNBC metastasis

Our studies as well as recent literature suggest that EZH2 plays a critical role in TNBC pathophysiology (Verma et al., 2018; Verma et al., 2020; Yomtoubian et al., 2020). TNBC is a highly heterogeneous group of cancers; where intrinsic subtype analysis suggests approximately 80% of TNBC belong to basal-like category (Burstein et al., 2015; Lehmann et al., 2011). The basal-like or basaloid TNBC (B-TNBC) is one of the most aggressive, therapy-resistant, and metastatic tumors (Volk-Draper et al., 2012). Therefore, we focus our entire studies on basal like TNBC subtype. Here, we sought to dissect the role of NC-EZH2 versus H3K27me3 in TNBC progression. We have established 4T-1based animal model that closely mimics basal like TNBC progression under preclinical setting (Maheshwari et al., 2017). To explore the individual role of NC-EZH2 versus H3K27me3 function, we made 4T-1 stable cells either overexpressing EZH2 catalytically inactive (–ΔSET) or catalytically hyperactive (Y641-F) protein (Figure 1A-1B). Following subcutaneous and orthotopic inoculation of control and genetically modified 4T1 cells into mice, we observe that Y641-F tumor bearing mice have smaller tumor compered to control tumor bearing mice, while EZH2 (–ΔSET) tumor bearing mice have no significant change in the tumor growth compere to control tumor bearing group (Figure 1A-1B and Figure 1C-1E and Supplementary Figure 1A) however, Y641-F tumor bearing mice lost significant (p<0.001) body weight compared to control, during tumor progression (Figure 1F). Significant body weight loss in Y641-F tumor bearing mice prompted us to check the metastasis status of the particular group. To understand the metastatic potential of individual group of cells, we adapted the same system but here we use Luc-tagged 4T-1 cells and investigate metastatic progression through sequential live animal imaging. As shown in Figure 1G-1H, Y641-F tumor bearing mice demonstrate a significant (p<0.01) increase in metastasis compared to control. Therefore, accelerated metastasis may be the reason for the marked weight loss of Y641-F tumor bearing mice. Altogether, *in- vivo* studies established that H3K27me3 but not NC-EZH2 protein regulates TNBC metastasis. Increased metastasis in Y641-F cells encouraged us to further explore the impact of H3K27me3 on TNBC cell migration and invasion. Here, we performed wound healing and trans-well chamber assay to assess the migration and invasion respectively. In wound healing assay (Figure 1I-1J), we detect Y641-F mutant cells have significantly (p<0.01) higher migratory potential as these cells are found to close the wound more rapidly than wild type cells. Similarly, in trans-well chamber assay, Y641-F cells also show significantly (p<0.01) higher invasive capabilities than that of wild type cells (Figure 1K-1L). Altogether, both *in vivo* and *in- vitro* studies suggest that the selective hyperactivation of H3K27me3 promotes TNBC migration, invasion and metastasis.

**Figure 1:**
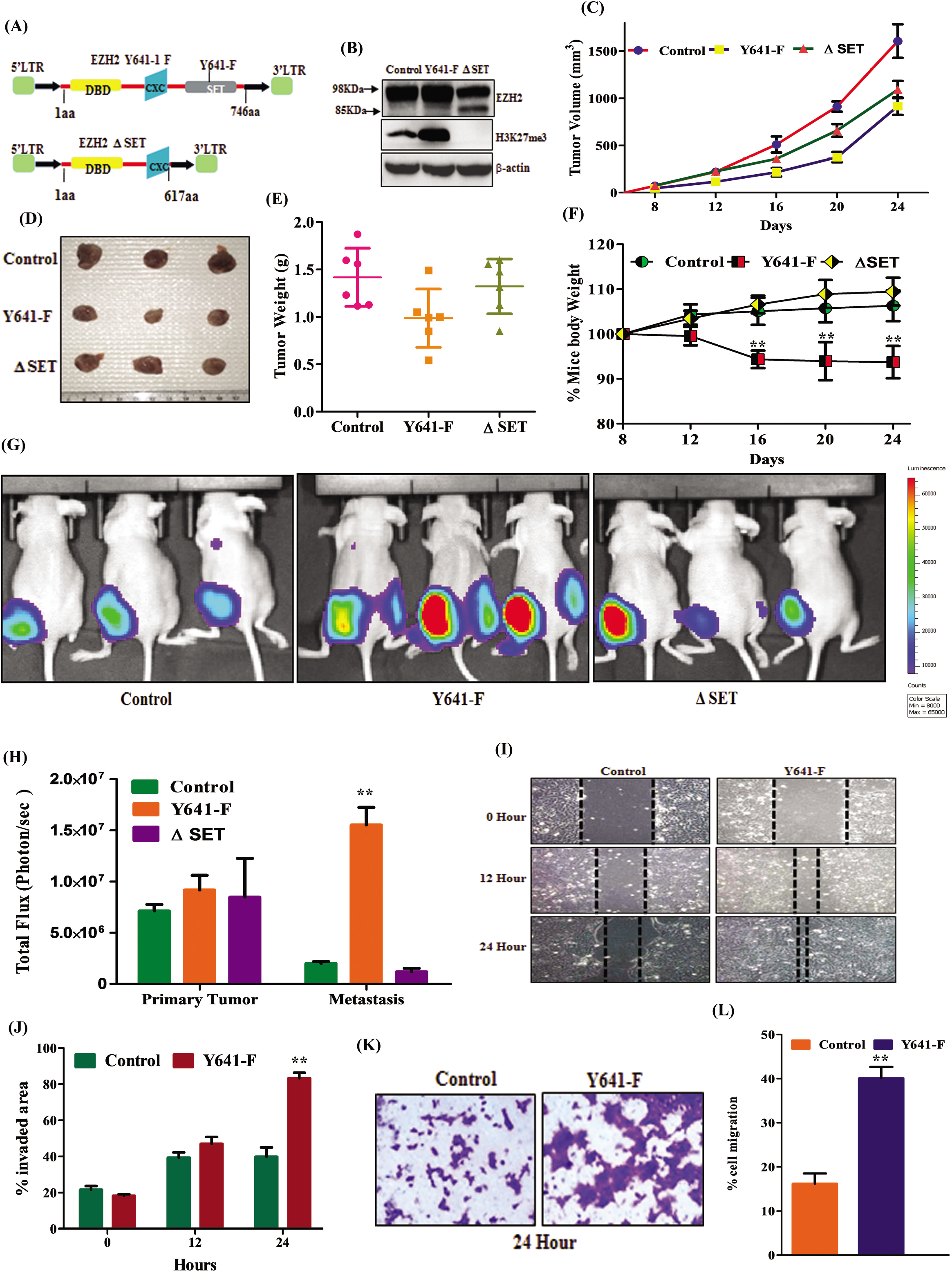
H3K27me3 but not NC-EZH2 protein promotes TNBC metastasis. (A) Schematic representation of EZH2 (Y641-F) and EZH2 –ΔSET retroviral overexpression constructs. (B) Immunoblot analysis for EZH2, H3K27me3, and β-actin expression in control, stable EZH2 (Y641-F), and EZH2 (ΔSET) overexpressed (OE) 4T-1 cells. (C) The control, EZH2 (Y641-F) OE and EZH2 ΔSET OE 4T-1 (1X10^6^) cells in 100 μl PBS were orthotopically inoculated in the right mammary fat pad of 4 to 6-week old female nude Crl: CD1-Foxn1nu mice (n=6) and allowed to grow for 25 days. Growth curve is shown in control, EZH2 (Y641-F) OE, and EZH2 –ΔSET OE; points are indicative of an average of tumor volume bars, (error bar, +/- SEM). (D) Representative images of harvested tumors belong to control, EZH2 (Y641-F) OE, and EZH2 –ΔSET OE bearing mice. (E) Average weight of harvested tumors, +/- SEM of Control, EZH2 (Y641-F) OE, and EZH2 –ΔSET OE groups depicted by graph. (F) The body weight curve is shown for control, EZH2 (Y641-F) OE, and EZH2 –ΔSET OE; points are indicative of average of body weight+/- SEM. *p-value <0.05, **p-value <0.001, Two Way ANOVA. (G) *In-vivo* bioluminescence monitoring of orthotopic control (left panel), EZH2 (Y641-F) OE 4T-1 Luc2-GFP (middle panel) and EZH2 – ΔSET OE (right panel) primary tumor and distant metastatic sites. The colour scale indicated the photon flux (photon/ sec) emitted from each group. (H) Quantitative bar graph representation of total photon flux calculated from the region of flux (ROI). **p- value, <0.001 student t-test. (I) Representative images of the wound healing assay to measure the migration ability of control and EZH2 (Y641-F) OE 4T-1 cells in time point manner (0 hour, 12 hour and 24 hour), magnification 10 X. (J) the quantitative analysis of wound healing assay.(Error bar, +/- SEM), **p-value, <0.001, Two way ANOVA.(K) representative images of the trans-well chamber migration assay to measure the invasion ability of control and EZH2 (Y641-F) OE 4T-1 cells at 24 hours. (L) The quantitative bar graphs of tans-well chamber migration assay of control and EZH2 (Y641-F) OE 4T-1 cells t 24 hours. ** P-value, <0.001, Student t-Test.

### Selective hyperactivation of H3K27me3 alters metastatic landscape of TNBC

To further validate the correlation between hyperactivation of H3K27me3 with rapid TNBC metastasis, we adapted three different strategies as described in (Figure 2A). Interestingly, in our uniquely designed three (mix single flank mammary fat pad, mix tail vein and individual double flank mammary fat pad) strategies, we observe that TNBC metastasis is robustly augmented upon H3K27me3 hyperactivation (Figure 2B-2C) compared to control. Next, we sought to determine the TNBC metastatic pattern under the influence of H3K27me3 hyperactivation. Fluorescence imaging analyses of harvested organs from double flank mammary fat pad inoculated (Control versus Y-641F) mice exhibit that control cells tend to metastasize mostly into the lungs whereas, Y-641F cells have a tendency to significantly metastasize into the spleen and liver but not to the lung (Figure 2 D). Representative fluorescent microscopic images and FACS analysis of single cells harvested from different metastatic organs of experimental mice clearly demonstrate the striking alteration of metastatic pattern observed with hyperactivation of H3K27me3 (Figure 2 E-G). Next, we sought to find out the correlation between H3K27me3 function and TNBC metastasis at basal state. We orthotopically inoculated WT 4T-1- Luc cells in the mammary fat pad of female nude mice (Figure 2H left panel). We isolated metastatic cells from different organs following mammary fat pad inoculation and after EZH2 expression analysis; we amazingly observe that splenic metastatic cells have highest expression of global H3K27me3 compared to cells isolated from primary tumors (Figure 2 H right panel). Notably, the splenic metastatic cells matched the phenotype with Y641-F cells at basal state. Further; we inoculated splenic isolated metastatic and primary tumor cells into the mammary fat pad again and evaluated the TNBC progression. As shown in (Figure 2 I-J), live animal imaging data clearly establish that splenic metastatic cell inoculation result in profound increase (p<0.01) of TNBC peritoneal metastasis compared to respective control. Further, splenic metastatic cell tumor bearing mice die significantly earlier than control (p< 0.01) (Figure 2K). Collectively, these series of *in- vivo* studies strongly suggest that the enhanced global trimethylation has a severe impact on TNBC peritoneal metastasis.

**Figure legends-2.**
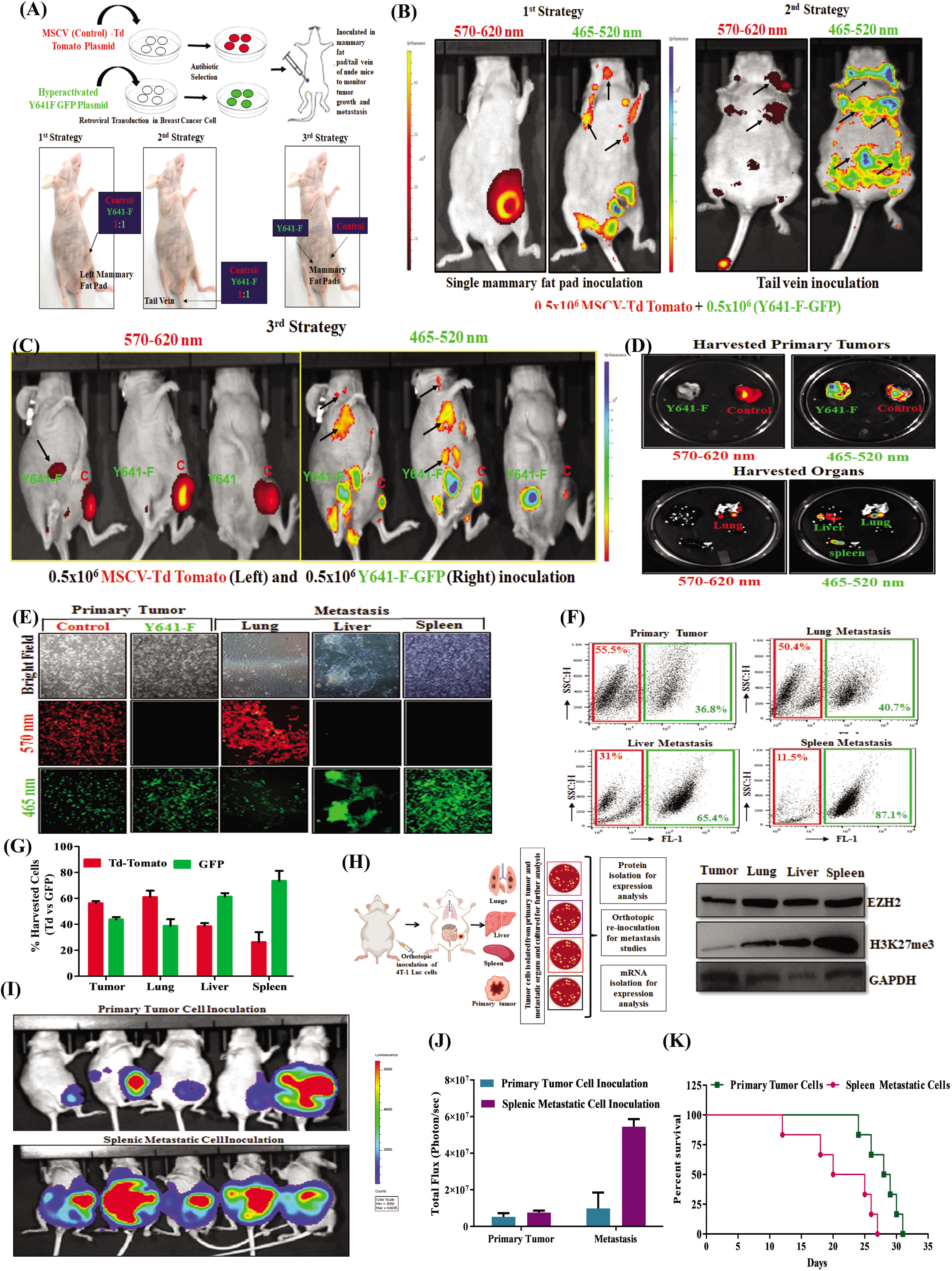
Hyper-activated H3K27me3 leads to TNBC splenic metastasis. (A) Schematic representation for generation of control Td-Tomato^+^ and hyper-activated H3K27me3 (Y641-F) GFP ^+^ 4T-1 cells by retroviral transduction (Upper panel). The schematic delineation of different strategies used in the *in-vivo* studies (Lower panel). First strategy: The Td-Tomato^+^ control and EZH2 (Y641-F) GFP^+^ cells were mixed in equal ratio (1:1). The 100μl of mixed cells (0.5 X 10 ^6^ Td- Tomato ^+^ 0.5 × 10 ^6^ Y641-F –GFP^+^) were orthotopically inoculated in the right mammary fat pad of female nude (n=5) (lower left panel). 2^nd^ strategy: The MSCV Td-Tomato^+^ control and EZH2 (Y641-F) GFP^+^ cells were mixed in equal ratio (1:1). The 100 μl of mixed cells (0.5 X 10 ^6^ Td-Tomato ^+^ 0.5 × 10 ^6^ Y641-F –GFP^+^) were inoculated *via* tail vein in the female nude mice (n=5) (lower middle panel). 3^rd^ strategy: the control (0.5 X 10 ^6^ Td-tomato ^+^) and EZH2 (0.5 × 10 ^6^ Y641 GFP ^+^) cells in 100 μl PBS were orthotopically inoculated in the right and left mammary fat pad of female nude mice (n=5) respectively (lower right panel). (B) 1^st^ strategy: the live fluorescence imaging of mixed orthotopic (control- Tomato ^+^ Y641-F –GFP^+^) primary tumors and distant metastatic sites (left panel). 2^nd^ strategy: the live fluorescence imaging of distinct metastatic sites in the mice group in which the mixed (control Td- tomato ^+^ +Y641 GFP ^+^) 4T-1 cells were inoculated *via* tail vein (Right panel). The excitation and emission wavelength: Td-Tomato- 570-620 nm and GFP-465-520 nm. (C) 3^rd^ strategy: the live fluorescence imaging for the identification of distant metastasis in the mice group developed the control (Td-Tomato ^+^) orthotopic tumor in the right and EZH2 Y641-F (GFP ^+^) orthotopic tumor in the left mammary fat pad. (D) The fluorescence imaging of harvested control (Td-Tomato ^+^) and EZH2 Y641-F (GFP ^+^) primary tumors at a specific wavelength (upper panel). The metastatic potential of control (Td-Tomato ^+^) and EZH2 Y641-F (GFP ^+^) cells were analysed by fluorescence imaging in the harvested organs at a specific wavelength (lower panel). (E) The fluorescence images of single-cell harvested from both tumors control (Td-Tomato ^+^) and EZH2 Y641-F (GFP^+^) and harvested organs (n=3). (F) Representative flow cytometry-derived scatter plots showing control (Td-Tomato ^+^) and EZH2 Y641-F (GFP ^+^) cells in primary tumors and metastatic organs harvested from mice (n=3). (G) The quantitative analysis of the percentage of Control (Td-Tomato^+^) and EZH2 Y641-F (GFP ^+^) cells in harvested primary tumors and metastatic organs (n=3) (Error bar, +/- SEM), p-value–ns, *p-value-< 0.01, **p-value, <0.001, Two way ANOVA. (H) The schematic representation of orthotopic 4T-1 cells inoculation, with isolation of primary tumor and metastatic cells from different organs (left panel), the immunoblot analysis to observed differential expression of EZH2, H3k27me3 in the basal 4T-1 cells harvested from primary tumor and metastatic organs. The GAPDH was used as loading control (right panel). (I) the *in-vivo* imaging of primary tumor inoculated cells and splenic metastatic cell inoculation bearing mice (n=5). The colour scale indicated the photon flux (photon/ sec) emitted from each group. (J) Quantitative bar graph representation of total photon flux calculated from the region of flux (ROI). *p- value, <0.001 student t-test.(K)The Kaplan-Meier survival curve of mammary tumor onset in the nude mice with inoculation of primary tumors cells (n=6) and spleen metastatic cells (n=6). *p- value, <0.05, log-rank test.

### Transcriptome analysis identified KRT14 as a new target of H3K27me3 in TNBC

H3K27me3 driven robust changes in TNBC metastatic signature encourages us to identify the mechanistic insight of the phenotype. To explore this idea, we perform differential Transcriptome analysis in control and Y641-F cells. The Venn diagram represents the common and differentially expressed genes between control and Y641-F (Figure 3A). Volcano plot represents significantly up and down regulated genes using the fold change cut off ≥2 between control and Y641-F cells (Figure 3B). This study reveals the upregulation of 92 and downregulation of 192 genes in Y641-F (p <0.05) group, compared to control. We extensively searched literature for metastasis related genes among 284 genes that were significantly altered in the Y641-F cells compered to control cells. The heat map represents the selected significantly up and downregulated genes which have a critical role in metastasis (p<0.05) (Figure 3C) (Coma et al., 2010; Gupta et al., 2007; Gurney et al., 2012; Jeng et al., 2020; Joosse et al., 2012; Lui et al., 2014; Nan et al., 2018; Oskarsson et al., 2011; Seachrist et al., 2017; Yin et al., 2010). Next, we carry out gene ontology for differentially expressed genes to identify the up and down regulated metastasis related KEGG pathways and most of the manually selected genes, we found common in the metastasis associated pathways (Supplementary Figure 2A, 2B and Supplementary Table S1, S2). Next, we validated the differential expression of the selected genes by the qRT-PCR between control and Y641-F cells. Likewise, the expression of *ATF-3, FZD5, ID2, ID3, KRT8, KRT14* and *KRT16* are found to be significantly (p<0.01) upregulated (Figure 3D), whereas, the expression of *FST, DOK7, TLE6, DEPTOR, PTPRUC* and *DLC-1* are significantly (p<0.01) downregulated in Y641-F cells as compared to control cells (Figure 3E). Next, to understand the contribution of these changes in ascertaining splenic metastasis in TNBC, we examined the mRNA expression of validated up and down regulated genes in the metastatic cells isolated from primary tumor, lungs, liver, and spleen. After having extensive validation, *KRT14* is found to be the most significantly (p<0.01) upregulated gene in the cells isolated from splenic metastasis as compared to primary tumor cells (Figure 3F), whereas, the mRNA expression of *DEPTOR* was found to be markedly (p<0.01) downregulated in the same (Figure 3G).

**Figure Legends 3.**
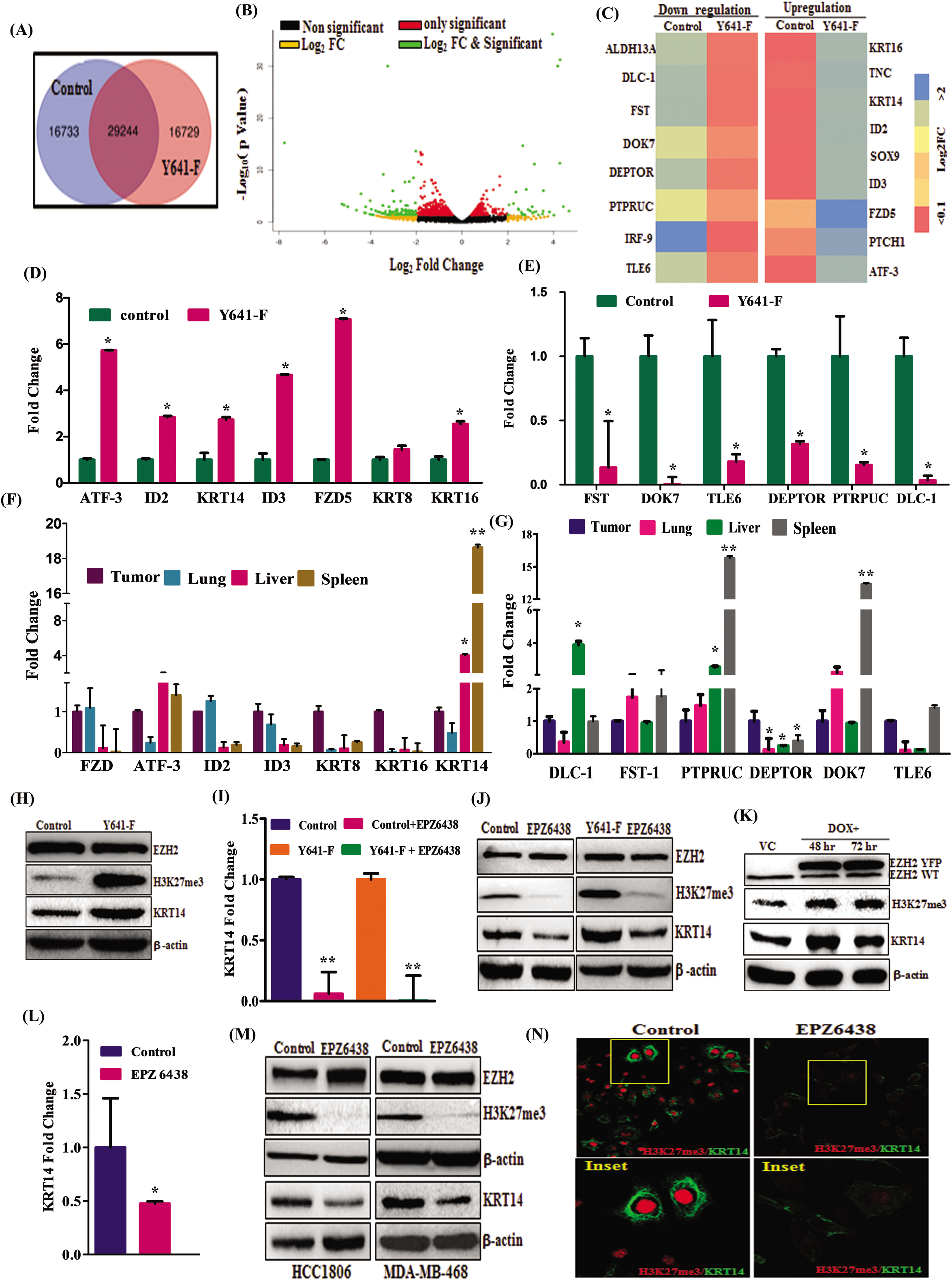
Hyper-activated H3K27me3 promotes TNBC metastasis by regulating KRT14. (A) Venn diagram summarizing the overlap between differentially expressed genes from RNA-Seq between control and EZH2 Y641-F cells. (B) Volcano plot represents the significant differential expression of genes in the form of Log2 (Fold change) of Fragment per Kilo base of transcript per Million mapped reads (FPKM) values between Control and EZH2 Y641-F. Data sets: Significantly up and downregulated genes in green (p<0.05). (C) Heatmap of signal intensity illustrating of differentially expressed metastasis related genes between control and EZH2 Y641-F cells using p<0.05 and fold change > 2.0 cut off. (D and E) Validation of up and downregulated genes identified in RNA-seq by real time-PCR between control and EZH2 Y641-F cells. Data points are average of triplicate readings of samples; error bars, ± S.D. *p<0.01, compared to control cells, student t-Test. (F and G) the differential expression of validated genes were analysed in the lungs, liver and spleen isolated metastatic cells with respect to primary tumor. Data points are average of triplicate readings of samples; error bars, ± S.D. *p<0.01, Student t-Test, compared to isolated tumor cells. (H) Control and EZH2 Y641-F cells were analysed for expression of EZH2, H3K27me3, KRT14 and β-actin by immunoblot. (I) the control and EZH2 Y641-F cells were treated with 10μM dose of EPZ6438 and performed q-PCR analysis for KRT14 expression. Data points are average of duplicate readings of samples; error bars, ± S.D. **p<0.001, compared to control cells.(J) The control and EZH2 Y641-F cells were treated with 10μM dose of EPZ6438 and analysed for expression of EZH2, H3k27me3, KRT14 and β-actin by immunoblot. (K)The vehicle control and Tet-ON EZH2-YFP OE HCC1806 cells were analysed for expression of EZH2 WT, EZH2-YFP, H3K27me3, KRT14 and β-actin by immunoblot.(L) The HCC1806 cells were treated either for vehicle control or 10μM dose of EPZ6438 for the analysis of the expression of KRT14 by RT-q PCR. Data points are average of duplicate readings of samples; error bars, ± S.D. **p<0.01, student t-Test compared to control cells.(M) The HCC1806 and MDAMB468 cells were either treated for vehicle or 10μM EPZ6438 to checked the expression of EZH2, H3K27me3, KRT14 and β-actin by immunoblot.(N) Confocal microscopy was performed for corresponding changes in global H3K27me3 level after vehicle or 10 μM EPZ6438 treatment for 24 hours. In similar setting, simultaneous expression of H3k27me3 and KRT14 was assessed by confocal microscopy.

Earlier reports (Au et al., 2013; Chen et al., 2020) regarding H3K27me3 mediated silencing of *DLC-1* and *DEPTOR* is supporting the authenticity of our current analysis. As our previous result suggested that the global H3K27me3 level was robustly high in the metastatic cells isolated from spleen compared to primary tumor cells (Figure 2H right panel). The KRT14 is the only gene which was found to be validated in both WT versus Y641-F cells and primary tumor versus splenic metastatic cells isolated from WT 4T-1 orthotropic inoculation (Figure 2H left panel). However, most of the other genes were failed to validate in the tumor cells versus splenic metastatic cells. Further, landmark study by Lehman et al., 2011 classified basal like TNBC on the basis of KRT gene family. Therefore, we concentrate our focus on H3K27me3 mediated KRT14 regulation and sought to validate our finding at protein level. Indeed, we observe KRT14 expression is robustly (p<0.01) upregulated in the Y641-F cells as compared to control and pharmacological inhibition of EZH2 catalytic activity by FDA approved drug tazemetostat (EPZ6438) also result in marked decrease in KRT14 expression at both mRNA and protein level (Figure 3H-3J). Further, we extend our validation experiments in two (HCC1806 and MDAMB468) different basal human TNBC cell lines. Inducible EZH2 gain of function in HCC1806 result in increase in KRT14 level (Figure 3K), while functional EZH2 inhibition by EPZ6438 in HCC1806 cells markedly (p< 0.01) downregulates the KRT14 mRNA expression (Figure 3L). Subsequently, the KRT14 protein expression was significantly downregulated in HCC1806 and MDAMB468 by EPZ6438 treatment. (Figure 3M). The downregulation of KRT14 were also observed in the EPZ6438 treated HCC1806 cells by confocal microscopy (Figure 3 N). Altogether, these findings clearly suggest that the methyltransferase activity of EZH2 promotes KRT14 expression during TNBC peritoneal metastasis.

### H3K27me3 enhances KRT14 transcription by attenuating binding of transcriptional repressor Sp1 to its promoter

Classically, H3K27me3 mark acts as a transcriptional repressor (Singh et al., 2020; Varambally et al., 2002). However, our data clearly indicate that it positively regulates KRT14 expression particularly in basal like TNBC subtype. To investigate whether H3K27me3 marks directly occupy the *KRT14* promoter that leads to transcription activation, we carry out the chromatin immunoprecipitation (ChIP) assay in 4T-1 and HCC1806 cells by using H3K27me3 and P-S5-RNA polymerase II specific antibody. To delineate the enrichment of H3K27me3 marks in the *KRT14* promoter, we design the walking primers in the sliding window of −3000 bases upstream and +2500 bases downstream of Transcription Start Site (TSS) by utilizing the Eukaryotic Promoter Database. The q-PCR analysis in the walking primer experiments reveal the strong enrichment of H3K27me3 marks at upstream of −0.2 kb and −0.5 Kb region in mouse 4T-1 cells, while in human HCC1806 cells the enrichment of H3K27me3 marks is selectively identified only in −0.2 kb region (p< 0.01). To determine whether H3K27me3 enrichment in the *KRT14* promoter is associated with transcription activation or repression, we perform the ChIP q-PCR for the transcription initiation specific RNA poll II C-terminal domain Ser-5 phosphorylation. Interestingly, the recruitment of p-Pol-II-S5 on the −0.2 kb region of *KRT14* promoter is observed, indicating the transcriptional activation in both 4T-1 and HCC1806 cells (Figure 4B-C). Similarly, we perform the ChIP q-PCR for *DLC1* gene as a positive control which is already known to be suppressed by H3K27me3. The q-PCR analysis of *DLC1* gene suggests the enrichment of H3K27me3 marks in the *DLC1* promoter. However, the recruitment of p-Pol-II-S5 is found to be minimal in the DLC-1 promoter, indicating the transcription repression of *DLC1* gene in 4T-1 cells (Supplementary Figure 3A and 3B) (Au et al., 2013). To further confirm the H3K27me3 mediated transcription activation of *KRT14,* we compare the enrichment of H3K27me3 in the control and Y641-F cells by ChIP-qPCR. Most likely, the robust (p<0.001) enrichment of H3k27me3 and p-Pol-II-S5 is found in Y641-F cells in comparison to the control cells. Robust (p<0.01) reduction of both H3K27me3 marks and p-Pol-II-S5 is identified in the *KRT14* promoter following EPZ6438 (10μM) treatment in HCC1806 cells, further establish the authenticity of the earlier observations (p< 0.01) (Figure D-F). Though H3K27me3 mark classically acts as a repressor, surprisingly, we find at least in our case, it promotes *KRT14* gene transcription.

**Figure legends: 4.**
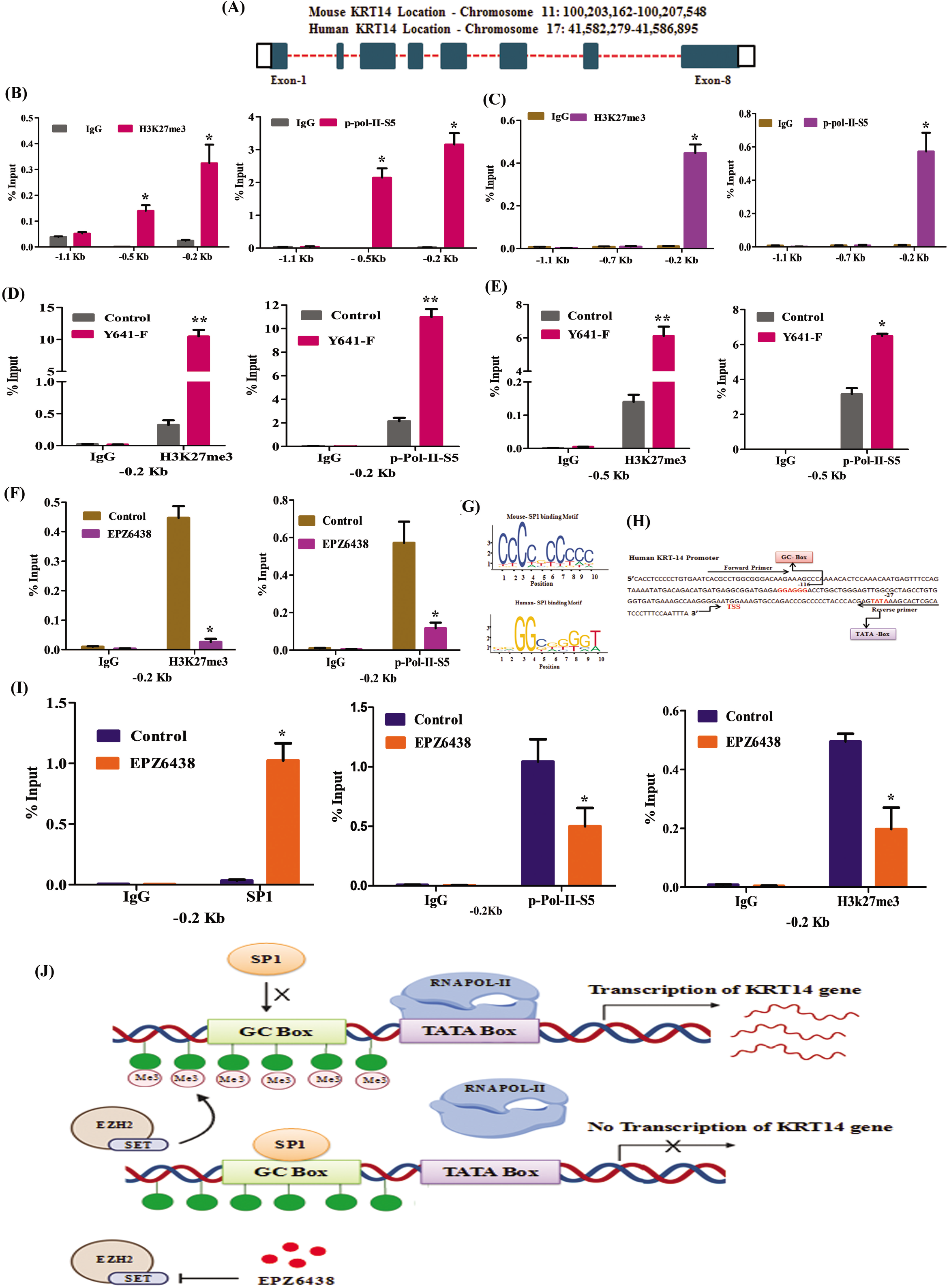
H3K27me3 enrichment in the KRT14 promoter is associated with active transcription. (A) Schema showing genomic location of KRT14 gene for both mouse and human. (B) ChIP was performed in 4T-1 mouse cells using anti-H3K27me3, p-Pol-II-S5 and IgG antibodies and then examined by real-time q-PCR using primer pairs targeting −1.5 Kb to +0.5 Kb of the *KRT14* gene (C) Same procedure for HCC1806 human cells as in B. (D and E) the differential fold change enrichment for H3K27me3 and p-Pol-II-S5 were observed in −0.2Kb and −0.5 Kb regions from TSS in control and EZH2-Y641 cells by ChIP q-PCR. (F) The differential fold change enrichment for H3K27me3 and p-Pol-II-S5 were analysed in −0.2Kb region from TSS in the HCC1806 cells treated either with vehicle control or 10μM EPZ6438.(G)Schema showing SP1 binding motif obtained from JASPAR database for mouse and human (Left panel).The human nucleotide sequence for *KRT14* promoter with TATA and GC-box. (H) ChIP q-PCR data showing the recruitment of SP1on the *KRT14* promoter upon EPZ6438 (10μM) treatment in HCC1806 cells (left panel). The differential enrichment of H3k27me3 (middle penal) and p-Pol-II-S5 (right panel) at *KRT14* promoter indentified by ChIP q-PCR. Results shown (A)-(H) are representative of two biological independent Experiments. Columns, an average of triplicate readings of samples; error bars, ± S.D. **, p <0.05 and **p value <0.001, student t-Test. (J) Illustrating showing that H3K27me3 inhibits the Sp1 binding in the promoter of *KRT14* gene and activate its transcription.

To gain further insight for H3K27me3 mediated KRT14 transcriptional activation, we search the transcription factors that regulate *KRT14* gene expression through TRRUST online software (www.grapedia.org)(Cai et al., 2012). We found that Sp1 act as a major transcriptional regulator of *KRT14.* The role of Sp1 has been highly characterized both as transcriptional activator and repressor (Hagen et al., 1994; Innocente and Lee, 2005). We analyse the presence of putative binding motif of Sp1 by employing publicly available transcription factor binding prediction software JASPAR (http://www.jaspar.genereg.net). The Sp1 usually binds to the GC-box of the promoter of the genes and regulates the transcription process. By utilizing the publicly available software Eukaryotic Promoter Database (epd.vital-it.ch,) we search the GC-box in the promoter of KRT14 to confirm the sequence of DNA binding motif of Sp1 (Figure 4G and H). We have identified the GC- box −116 base upstream of TSS in *KRT14* promoter. Interestingly, the enrichment of H3k27me3 marks and p-ser-5-Poll have also been recognized in the same region. To examine whether H3K27me3 regulates the recruitment of Sp1 in the GC-box, we first confirm the binding of Sp1 in GC box of *KRT14* promoter. We performed the ChIP q-PCR in the Sp1 immunoprecipitated DNA in HCC1806 control and EPZ6438 (10μM) treated cells. Interestingly, the recruitment of Sp1 is found to be robustly (p<0.01) up regulated in the EPZ6438 cells, compared to control, while, the H3K27me3 and p-Pol-II-S5 enrichment in the KRT14 promoter is significantly (p<0.01) down regulated in EPZ6438 treated cells as compared to the control cell (Figure 4I and J). All together, these finding suggest that the H3K27me3 may compact the GC box region in *KRT14* promoter to inhibit the binding of Sp1 in the GC box and concede RNA polymerase II to initiate the transcription of *KRT14* gene.

### Genetic ablation of KRT14 impairs splenic metastasis in TNBC

H3K27me3 mediated selective KRT14 upregulation in TNBC peritoneal metastasis encouraged us to explore the role of KRT14 in TNBC migration, invasion, metastasis. First, we knock down the KRT14 in 4T-1 (Y641-F) and HCC1806 cells by using the two-specific shRNA against KRT14 mRNA and validated their efficacy (Figure 5 A and B). In wound healing assay, we find KRT14 KD significantly (p<0.01) inhibits closure of wound as compared to control (Figure 5 C and F). Similarly, in the trans-well chamber assay, we observe KRT14 KD cells have significantly (p<0.01) lower invasion capabilities than control cells (Figure 5G and J). To determine the role of KRT14 in TNBC splenic metastasis, we perform the orthotopic single mammary fat pad mixture experiment where (Y641-F) control Td-Tomato^+^ and (Y641-F) KRT14 KD GFP^+^ cells are equally mixed and inoculated in one of the mammary fat pads of mice (n=5). After 25 days, we harvested spleen and isolated the single cells from the spleen and allowed them to grow for 3 days with 6-TG selection. As shown in fluorescence microscopy pictures (Figure 5K and L), majority of metastatic cells migrated towards spleen are found to be control Td-Tomato^+^ cells as compared to GFP^+^ KRT14 KD cells. Next, cells were trypsinized and analysed by FACS to quantitatively analyse the percentage of Control and KRT14 KD cells metastasized to spleen. Our FACS analysis clearly suggests that the percentage of migrated metastatic control Td-tomato^+^ cells are significantly (p<0.01) higher in the spleen as compared to metastaticY641-F GFP^+^ cells (Figure 5M and 5N). Altogether, these studies conclusively demonstrate that loss of KRT14 expression reduces TNBC cell migration and invasion capabilities and markedly hinders TNBC splenic metastasis.

**Figure Legends 5:**
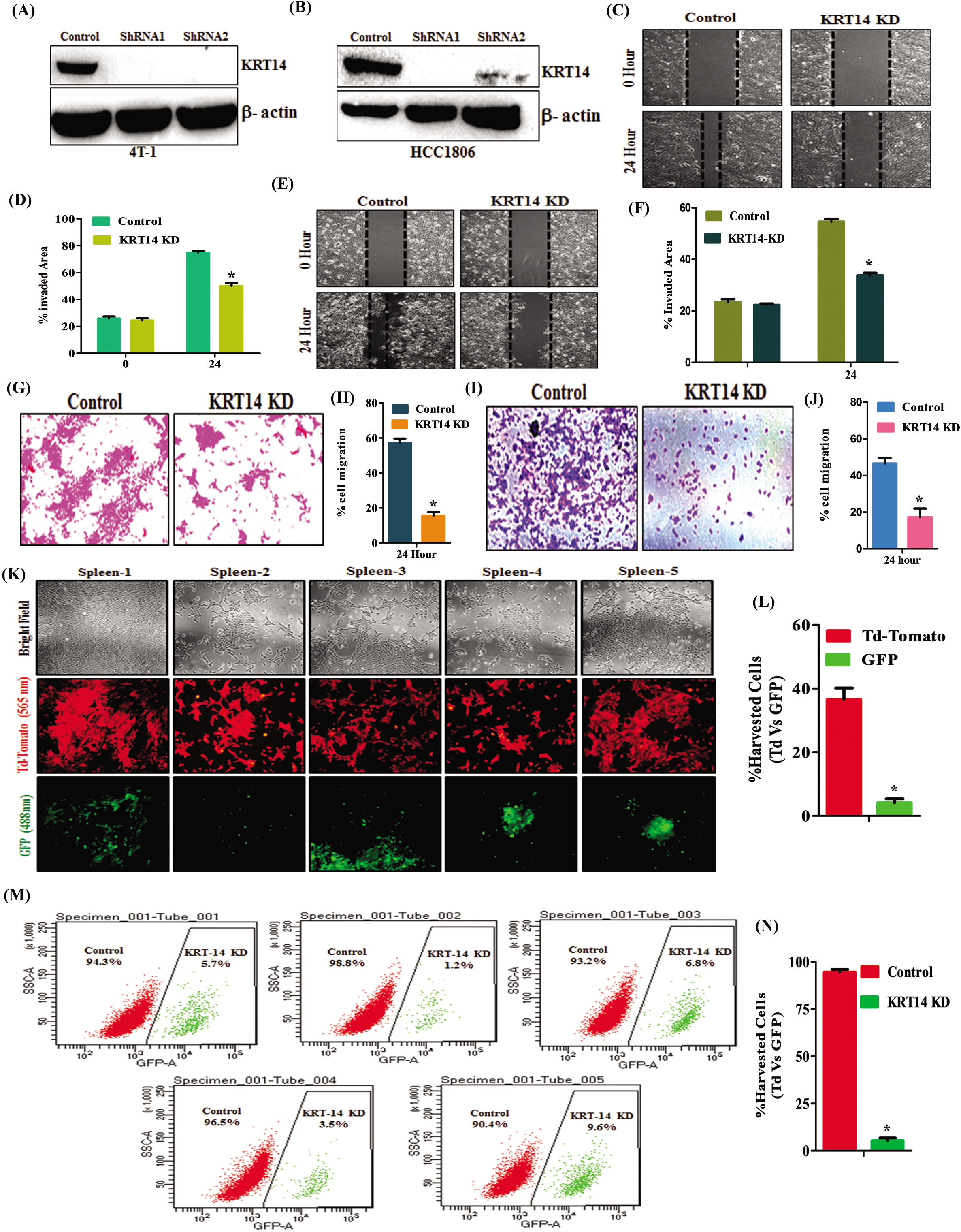
Genetic knock down of KRT14 inhibits splenic metastasis. (A and B)The KRT14 knock down was confirmed through immunoblot by using two different shRNA in 4T-1 and HCC1806 cells. The β-actin used as a loading control. (C) Representative images of the wound healing assay to measure the migration ability of control and KRT14 KD in 4T-1 cells in 24 hours, magnification 10 X. (D) The quantitative analysis of wound healing assay.(Error bar, +/- SEM), *p-value, <0.01,Two Way ANOVA.(E)Same wound healing process for HCC1806 cells as in (C). (F) The quantitative analysis of wound healing assay.(Error bar, +/- SEM), *p-value, <0.01,Two Way ANOVA.(G) Representative images of the trans-well chamber migration assay to measure the invasion ability of control and KRT14 KD 4T-1 cells at 24 hours. (H) The quantitative bar graphs of tans-well chamber migration assay of control and KRT14 KD 4T-1 cells at 24 hours. * P-value, <0.01, Student t-Test. (I) the same cell migration assay was performed in HCC1806 as in (G). (J)The quantitative analysis of trans-well assay, (Error bar, +/- SEM), **p-value, <0.01, Student t-Test.(K) Fluorescence imaging of metastatic cells harvested from spleen (n=5).(l) The Td-Tomato fluorescence is for control cells and GFP fluorescence is for KRT14 KD (Left panel).The excitation and emission wavelength: Td-Tomato- 570-620 nm and GFP-465-520 nm. The quantitative analysis for the metastatic control (Td-Tomato ^+)^ and KRT14 KD (GFP^+^) cells harvested from spleen (n=5). (M) Representative flow cytometry-derived scatter plots showing control (Td-Tomato ^+^) and EZH2 Y641-F (GFP^+^) metastatic cells harvested from spleen (n=5). (N) The quantitative analysis of the percentage of Control (Td-Tomato^+^) and KRT14 KD (GFP ^+^) metastatic cells harvested from spleen (n=5) (Error bar, +/- SEM),*p-value, <0.001, Two way ANOVA.

### Inhibition of EZH2 impairs TNBC peritoneal metastasis

We wish to explore the clinical relevance of our finding further and trying to finally understand that whether EZH2 functional inhibition rescues the *in- vivo* phenotype. In this endeavour, we first confirm EZH2 knockdown in 4T-1 cells through immunoblot (Figure 6 A). Indeed, EZH2 knockdown cells display significantly (p<0.01) less migratory (Figure 6B and C) and invasive (Figure 6 D and E) potential than its respective controls (p<0.01). Further, the serial bioluminescence imaging and analysis of harvested organs from control EZH2 KD tumor bearing mice reveal that the depletion of EZH2 robustly (p<0.001) restricts the peritoneal metastasis as compared to respective control (Figure 6F and H). Also, compared to control, EZH2 KD significantly (p<0.001) increases mice survival as shown in (Figure 6I). As EZH2 inhibitor EPZ6438 (tazemetostat) recently received FDA approval for sarcoma treatment, we readily evaluated its potential for the inhibition of TNBC migration, invasion and metastasis (Hoy, 2020; Rothbart and Baylin, 2020). As observed in (Figure 6J - M), EPZ6438 treatment result in significant (p<0.01) inhibition of TNBC migration and invasion, compared to their respective controls. Consistent with the genetic depletion of EZH2, we observe a dramatic (p <0.01) decrease of peritoneal metastasis following daily oral administration of EPZ6438 (250mg/kg) though primary tumor burden remains same in both control and treatment group (Figure 6N-6P and Supplementary Figure 4A). Moreover, EPZ6438 treatment results in significant (p<0.01) increase in mice survival as compared to vehicle treated group (Figure 6Q). Interestingly, the TCGA data mining of breast cancer patients shows the positive correlation between EZH2 and KRT14 expression only in the basal (TNBC) subtype compared with other counterparts (Figure 6R-6S). In summary, our *in vitro* and *in vivo* data together suggest that FDA approved drug EPZ6438 (H3K27me3 inhibitor) has immense potential to be developed as a targeted therapy against TNBC metastasis.

**Figure: 6.**
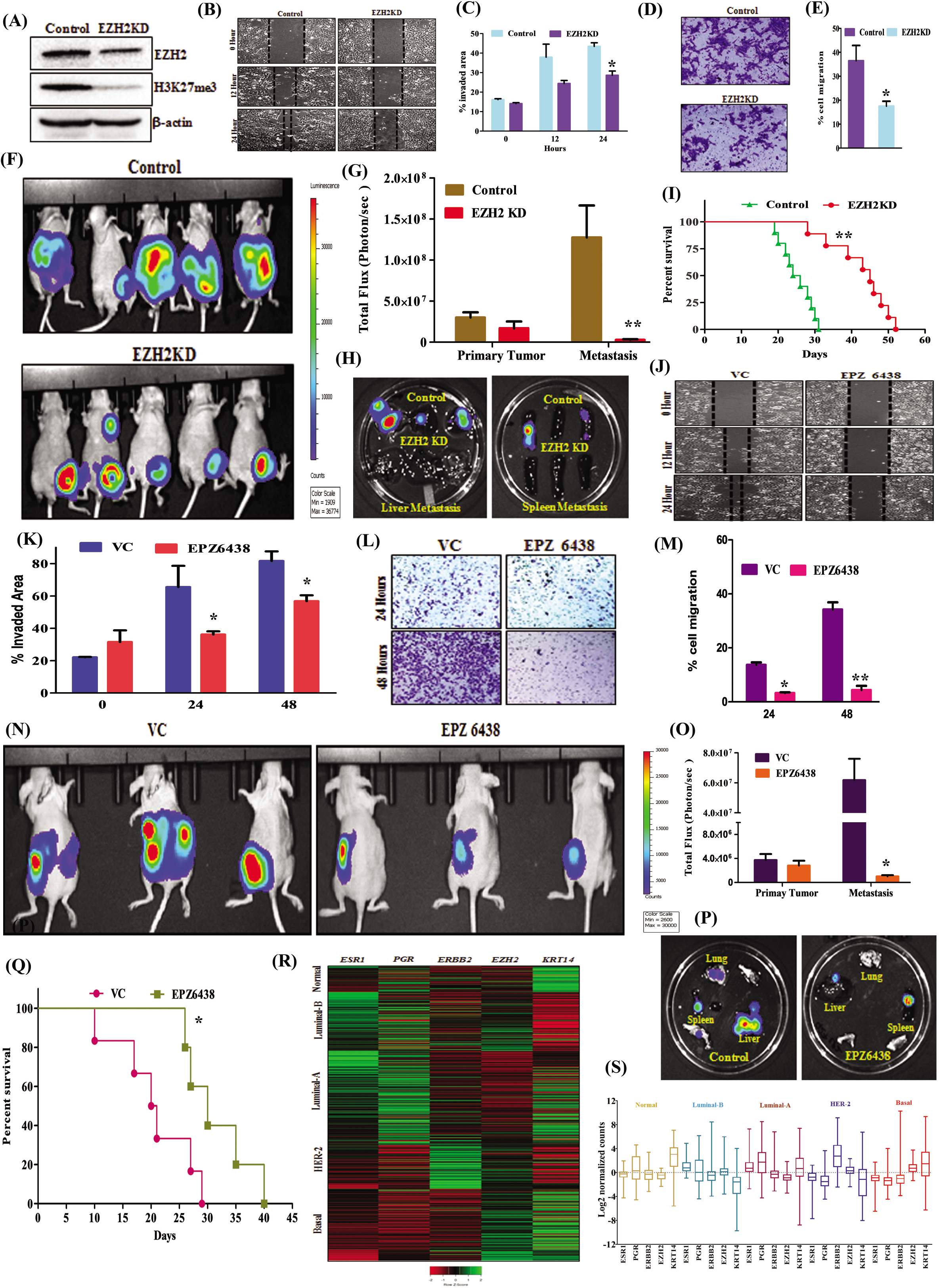
Inhibition of EZH2 methyltransferase activity reduces the peritoneal metastasis in TNBC. A) Stable control and EZH2 knockdown (KD) 4T-1cells were analyzed for expression of EZH2, H3K27me3 and β-actin by immunoblot. (B)Representative images of the wound healing assay to analyse the migration ability of control and EZH2 KD 4T-1 cells in time point manner (0 hour, 12 hour and 24 hour), magnification 10 X. (C) The quantitative analysis of wound healing assay. (Error bar, +/- SEM), *p-value, <0.01, Two way ANOVA.(D) Representative pictures of the trans-well chamber migration assay to analyse the invasion ability of control and EZH2 KD at 24 hours. (E) The quantitative bar graphs of tans-well chamber migration assay of control and EZH2 KD 4T-1 cells at 24 hours. * P-value, <0.01, Student t-Test (F) *In-vivo* bioluminescence imaging of orthotopic control (Upper penal), EZH2 KD 4T-1 Luc2- GFP (Down panel) primary tumor and distant metastatic sites. The colour scale indicated the photon flux (photon/ sec) emitted from each group. (G) Quantitative bar graph representation of total photon flux calculated from the region of flux (ROI). **p- value, <0.001 student t-test.(H) Bioluminescence analysis of control and EZH2 KD metastatic cells migrated in the liver and spleen.(I) The Kaplan-Meier survival curve of mammary tumor onset in the nude mice with inoculation of control (n=10) and EZH2 KD cells (n=10).**p- value, <0.001, log-rank test.(J) Representative pictures of wound healing assay to observe the migration ability of EZH2Y641-F cells under the treatment of EPZ6438 (10 μM). (K) The quantitative analysis of wound healing assay, (Error bar, +/- SEM), *p-value, <0.01, Two way ANOVA. (L) Representative images of trans- well chamber assay to analyse the invasion ability of control and EPZ6438 (10 μM) treated cells. (M)The quantitative analysis of wound healing assay * P-value, <0.01 and ** p value <0.001, Student t-Test.(N) The EZH2 (Y641-F) (1X10^6^) cells in 100 μl PBS were orthotopically inoculated in the right mammary fat pad of 4-to 6-week-old female nude (n=10 each group) and allowed to grow for 25 days. After 1 week mice were divided in to vehicle and treatment groups (n=5) each group. The EPZ6438 (200mg/Kg) dose was administered to mice once in a day by oral route. *In-vivo* bioluminescence monitoring of orthotopic nude mice, either treated with vehicle control (Y641-F) (n=5) or EPZ6438. The colour scale indicated the photon flux (photon/ sec) emitted from each group. (O) Quantitative bar graph representation of total photon flux calculated from the region of flux (ROI).*p- value, <0.05 student t-test. (P) The bioluminescence analysis of metastatic signal in the organs harvested from control (Y641-F) and EPZ6438 treated mice.(Q)The Kaplan-Meier survival curve of nude mice, either treated with vehicle control (n=5) or EPZ6438 (5mg/Kg) (n=5).*p- value, <0.05, log-rank test. (R) Analysis of EZH2 and KRT14 transcript levels on the basis of ESR1, PGR, ERBB2 expression in different PAM50 subtypes represented in a heat map. Data is retrieve from Breast Cancer (Yau2010) data set, publicly accessible via XENA USSC Cancer Genome Browser.( S) Representative of heat map in the form of Whiskers plot for Breast Cancer (Yau2010) PAM50 subtypes, Normal (Brown) n=66, ESRI s.d.=1.01, mean=-0.26, PGR s.d.= 2.35, mean= 0.474, ERBB2s.d.=1.41, mean=-0.212, EZH2 s.d.= 0.80, mean= −.505 and KRT14 s.d=2.33, mean=2.76. Luminal B (sky blue) n=139, ESR1 s.d.=0.39, mean=0.071, PGR, s.d.= 2.44, mean=0.155, ERBB2 s.d.=1.36,mean=-0.420, EZH2 s.d. =1.20, mean=0.366, KRT14 s.d=2.47, mean=-2.28.Luminal A n=222, ESR1 s.d.=0.45, mean= −0.301, PGR s.d.= 2.27, mean=1.26, ERBB2 s.d.=1.43, mean=-0.195, EZH2 s.d.=0.725, mean=-0647 and KRT14 s.d.= 2.80, mean=0.752. HER+ n=102, ESR1 s.d. = 1.70, mean=-1.842, PGR s.d. =1.56, mean=-1.48, ERBB2, s.d. =2.59, mean=3.29, EZH2 s.d. =0.78, mean=0.303 and KRT14 s.d. =2.79, mean=-1.906. Basal n=170, ESR1s.d. =1.80, mean=-2.76, PGR s.d. =0.791, mean=-1.54, ERBB2 s.d. =2.50, mean=-1.18, EZH2 s.d. =0.98, mean=1.97 and KRT14 s.d. =3.42, mean=1.45. Whiskers for the plot signify s.d. and the bar denotes the mean for each subtypes.

## Discussion

In the current study, we set out to dissect H3K27me3 versus NC-EZH2 in basal like TNBC growth and progression and discover that H3K27me3 is the key for TNBC peritoneal metastasis. Transcriptome analyses and subsequent validation lead us to ascertain KRT14 as a new target of H3K27me3 and most astonishingly, we observe increased H3K27 tri-methylation mark for transcriptional activation of KRT14 instead of H3K27me3 mediated classical transcription repression. Loss of EZH2 or KRT14 or H3K27me3 function results in robust inhibition of TNBC peritoneal metastasis, particularly splenic metastasis.

The advancement in the high throughput techniques enlightens EZH2 as a TNBC specific target. Recently, performed a bioinformatics analysis of a 2,000 patients in the breast cancer cohort and identified that TNBC patients have shown the enhanced expression of EZH2 with over all poor survival rate (Yomtoubian et al., 2020). Moreover, several human tissue microarray studies correlated EZH2 with poor prognosis in TNBC (Guo et al., 2016; Hussein et al., 2012; Kleer et al., 2003). In support of all the pre-existing information, our studies further prove that particularly in basal like TNBC subtype, which covers 80% of the whole group (Lehmann et al., 2011), EZH2 activity is a critical driver for TNBC progression.

Moreover, EZH2 overexpression in breast cancer does not always correlate with increased expression of global H3K27me3 (Hirukawa et al., 2018; Holm et al., 2012; Ma et al., 2020). It is inadequately established whether TNBC with distinct levels of EZH2 protein and its catalytic function (H3K27me3) belong to the same or different biological behaviour. In this context, we observe that H3K27me3 but not NC-EZH2 protein plays a pivotal role in modulating metastasis at least in case of basal like TNBC which is greatly supported by recent work of Mittal group (Yomtoubian et al., 2020). Peritoneal metastasis is a classical signature of TNBC metastasis in patients which is largely missing in most of the earlier preclinical studies as route of tumor cell inoculation is not always orthotopic, rather tail vein or intra-cardiac (Compérat et al., 2007; Vishnoi et al., 2019). Further, it has been shown that peritoneal metastasis associated with poor survival among the other distant metastatic sites of breast cancer (Beniey, 2019; Bertozzi et al., 2015; Flanagan et al., 2018). Therefore, our observation regarding H3K27me3 dependent TNBC splenic metastasis actually displays the real clinical scenario and its further inhibition has tremendous translational impact in TNBC pathophysiology. Previously, an elegant study showed the impact of epigenetic reprogramming in the progression of pancreatic ductal adenocarcinoma (PDAC) and identified the substantial difference in the enrichment of repressive (H3K9me3) and activation (H3K27ac and H3K36me) marks between peritoneal and distant metastasis (McDonald et al., 2017). Similarly, we also identified the massive epigenetic reprogramming in the expression of global H3K27me3 between the isolated cells from primary tumor versus cells from peritoneal metastatic organs like as spleen and liver. Further, we observe isolated H3K27me3 enriched spleen metastatic cells have even severe disease progression and fatality in mice indicating the critical correlation between hyperactive H3K27me3 function and TNBC progression and its overall poor survival.

Our transcriptome analysis of H3K27me3 hyper-activated cells not only identifies cytokeratin 14 (KRT14) as a new target of H3K27me3 in TNBC but also establishes a unique correlation between induction of H3K27me3 with upregulation of gene *(KRT14)* expression. Usually, EZH2 mediated silencing of genes is the fundamental mechanism of the PRC2 complex (Schwartz and Pirrotta, 2007; Varambally et al., 2002). In spite of its classical suppressive function, Majewski group first documented enrichment of H3K27me3 in the transcriptionally activated genes (Young et al., 2011). In correlation with our finding, H3K36me3 and H3K4me2 have shown to play contrasting impact on gene expression depending on their differential distribution in chromatin landscape (Kolasinska-Zwierz et al., 2009; Pekowska et al., 2010). The evolution in the ChIP-Seq pipeline revealed that the chromatin landscape is highly dynamic and the distribution of epigenetic marks is not universal. Indeed, the changes in the classical distribution pattern of any epigenetic signature may change the global network of gene transcription (Hon et al., 2012; Kolasinska-Zwierz et al., 2009; Pekowska et al., 2010; Young et al., 2011). Further extension with the concept, here we provide new insight that recruitment of transcription factor can be controlled by epigenetic modulator as H3K27me3 promotes *KRT14* transcription by inhibiting the binding of transcription factor Sp1 to its promoter in TNBC cells. Besides novel epigenetic regulation of KRT14 expression, another important aspect of our study is the critical positioning of KRT14 as a potent metastatic regulator in TNBC. So far, it is well known for an intrinsic molecular marker for basal like TNBC (Burstein et al., 2015; Lehmann et al., 2011) but our studies demonstrate its functional significance in TNBC pathophysiology. In support of our observation, several reports have suggested that the KRT14^+^ cells have metastatic advantage in compared to KRT14^-^ cells in TNBC heterogeneous population and regulate the function of several genes involved in different stages of metastatic cascade. (Cheung et al., 2016; Granit et al., 2018; Yong et al., 2020). Further, KRT14 acts as a stem cell for the natural or injury induced bladder regeneration and typically lead the origin of bladder cancer (Papafotiou et al., 2016).

Existing literature indicate that role of EZH2 in tumor growth and progression is highly context dependent and to some extent it exerts completely opposite effect in different types of cancers. Our recent studies in colon cancer, as well as an elegant study by Gonzalez et.al, (Gonzalez et al., 2014; Singh et al., 2020) in breast cancer exhibit loss of EZH2 results in primary tumor growth inhibition, whereas, our current observation in TNBC shows no significant impact of EZH2 in primary tumor growth. Consistent with our anti-metastatic effect of EZH2, recently (Hirukawa et al., 2018; Yomtoubian et al., 2020) tested the effect of EZH2 HMT inhibitor (GSK-126) against TNBC and luminal B breast cancer respectively and witness the robust anti-metastatic potential of EZH2 inhibitor. Context dependency is profound even in terms of EZH2 dependent gene regulation as our TCGA data mining in the PAM50 subtypes of breast cancer cohort display direct positive correlation between EZH2 and KRT14 expression selectively in basal breast cancer subtype (TNBC) among all breast cancer subtypes. Therefore, clinician should consider EZH2 therapeutic window very carefully as EPZ6438 (EZH2 inhibitor) or Tazemetostat recently received FDA approval against sarcoma (Hoy, 2020; Rothbart and Baylin, 2020).

In conclusion, our results reveal a new mechanism whereby EZH2 catalytic activity enhances the KRT14 expression in the basal like TNBC subtype. Mechanistically, we have shown instead of classical suppression; how H3K27me3 can promote gene expression like KRT14 transcription in order to regulate TNBC peritoneal metastasis. Finally, we identify the EZH2 inhibitor drug Tazemetostat (EPZ6438) can be a promising therapeutic candidate against the most aggressive TNBC subtype, where targeted therapy is still an enigma.

## Availability of Data and Materials

All data needed to evaluate the conclusions in the paper are present in the paper and/or in the Supplementary Materials. Additional data related to this paper will be made available upon reasonable request.

## Acknowledgments

We sincerely acknowledge the excellent technical help of Mr. A. L. Vishwakarma of SAIF for the Flow Cytometry studies; Ms Reema of Electron Microscopy unit for Confocal Imaging. We express our deepest gratitude to Dr. SK Rath and Mr. Navadayam for providing the Imaging facility. Research of all the authors’ laboratories was supported by CSIR-CDRI Institutional Fund (MLP019) and Fellowship grants from CSIR, DBT, and UGC. D.D acknowledges grant support from DST (EMR/2016/006935), DBT (BT/AIR0568/PACE-15/18) and ICMR (2019-1350). Institutional (CSIR-CDRI) communication number for this article is 135.

## Author Contributions

AV involved in study designing, performed experiments and wrote the draft manuscript. AS helped in carrying out *in vivo* studies. AKS, MAN, KKS provided active support for carrying out various *in vitro* and *in vivo* experiments. AS supplied critical reagents and provided intellectual input. DD conceived the idea, designed experiments, analysed data, wrote the manuscript and provided overall supervision. All authors read and approved the final manuscript.

## References

Au, S.L.-K., Wong, C.C.-L., Lee, J.M.-F., Wong, C.-M., and Ng, I.O.-L.J.P.o. (2013). EZH2-mediated H3K27me3 is involved in epigenetic repression of deleted in liver cancer 1 in human cancers. 8, e68226.

Beniey, M.J.C. (2019). Peritoneal metastases from breast cancer: a scoping review. 11.

Bertozzi, S., Londero, A.P., Cedolini, C., Uzzau, A., Seriau, L., Bernardi, S., Bacchetti, S., Pasqual, E.M., and Risaliti, A.J.S. (2015). Prevalence, risk factors, and prognosis of peritoneal metastasis from breast cancer. 4, 688.

Bray, F., Ferlay, J., Soerjomataram, I., Siegel, R.L., Torre, L.A., and Jemal, A.J.C.a.c.j.f.c. (2018). Global cancer statistics 2018: GLOBOCAN estimates of incidence and mortality worldwide for 36 cancers in 185 countries. 68, 394–424.

Burstein, M.D., Tsimelzon, A., Poage, G.M., Covington, K.R., Contreras, A., Fuqua, S.A., Savage, M.I., Osborne, C.K., Hilsenbeck, S.G., and Chang, J.C.J.C.C.R. (2015). Comprehensive genomic analysis identifies novel subtypes and targets of triple-negative breast cancer. 21, 1688–1698.

Cai, B.-H., Hsu, P.-C., Hsin, l.-L., Chao, C.-F., Lu, M.-H., Lin, H.-C., Chiou, S.-H., Tao, P.-L., and Chen, J.-Y.J.P.O. (2012). p53 acts as a co-repressor to regulate keratin 14 expression during epidermal cell differentiation. 7, e41742.

Chaffer, C.L., and Weinberg, R.A.J.S. (2011). A perspective on cancer cell metastasis. 331, 1559–1564.

Cheang, M.C., Voduc, D., Bajdik, C., Leung, S., McKinney, S., Chia, S.K., Perou, C.M., and Nielsen, T.O.J.C.c.r. (2008). Basal-like breast cancer defined by five biomarkers has superior prognostic value than triple-negative phenotype. 14, 1368–1376.

Chen, S.-q., Li, J.-q., Wang, X.-q., Lei, W.-j., Li, H., Wan, J., Hu, Z., Zou, Y.-w., Wu, X.-y., and Niu, H.-x.J.C.D. (2020). EZH2-inhibitor DZNep enhances apoptosis of renal tubular epithelial cells in presence and absence of cisplatin. 15, 1–11.

Cheung, K.J., Padmanaban, V., Silvestri, V., Schipper, K., Cohen, J.D., Fairchild, A.N., Gorin, M.A., Verdone, J.E., Pienta, K.J., and Bader, J.S.J.P.o.t.N.A.o.S. (2016). Polyclonal breast cancer metastases arise from collective dissemination of keratin 14-expressing tumor cell clusters. 113, E854–E863.

Coma, S., Amin, D.N., Shimizu, A., Lasorella, A., Iavarone, A., and Klagsbrun, M. (2010). Id2 promotes tumor cell migration and invasion through transcriptional repression of semaphorin 3F. Cancer Res 70, 3823–3832.

Compérat, E., Bardier-Dupas, A., Camparo, P., Capron, F., Charlotte, F.J.A.o.p., and medicine, I. (2007). Splenic metastases: clinicopathologic presentation, differential diagnosis, and pathogenesis. 131, 965–969.

Dent, R., Trudeau, M., Pritchard, K.I., Hanna, W.M., Kahn, H.K., Sawka, C.A., Lickley, L.A., Rawlinson, E., Sun, P., and Narod, S.A.J.C.c.r. (2007). Triple-negative breast cancer: clinical features and patterns of recurrence. 13, 4429–4434.

Flanagan, M., Solon, J., Chang, K., Deady, S., Moran, B., Cahill, R., Shields, C., and Mulsow, J.J.E.J.o.S.O. (2018). Peritoneal metastases from extra-abdominal cancer–a population-based study. 44, 1811–1817.

Gonzalez, M.E., Moore, H.M., Li, X., Toy, K.A., Huang, W., Sabel, M.S., Kidwell, K.M., and Kleer, C.G.J.P.o.t.N.A.o.S. (2014). EZH2 expands breast stem cells through activation of NOTCH1 signaling. 111, 3098–3103.

Granit, R.Z., Masury, H., Condiotti, R., Fixler, Y., Gabai, Y., Glikman, T., Dalin, S., Winter, E., Nevo, Y., and Carmon, E.J.C.r. (2018). Regulation of cellular heterogeneity and rates of symmetric and asymmetric divisions in triple-negative breast cancer. 24, 3237–3250.

Guo, S., Li, X., Rohr, J., Wang, Y., Ma, S., Chen, P., and Wang, Z.J.D.P. (2016). EZH2 overexpression in different immunophenotypes of breast carcinoma and association with clinicopathologic features. 11, 1–9.

Gupta, G.P., and Massagué, JJ.C. (2006). Cancer metastasis: building a framework. 127, 679–695.

Gupta, G.P., Perk, J., Acharyya, S., de Candia, P., Mittal, V., Todorova-Manova, K., Gerald, W.L., Brogi, E., Benezra, R., and Massague, J. (2007). ID genes mediate tumor reinitiation during breast cancer lung metastasis. Proc Natl Acad Sci U S A 104, 19506–19511.

Gurney, A., Axelrod, F., Bond, C.J., Cain, J., Chartier, C., Donigan, L, Fischer, M., Chaudhari, A., Ji, M., Kapoun, A.M., et al. (2012). Wnt pathway inhibition via the targeting of Frizzled receptors results in decreased growth and tumorigenicity of human tumors. Proc Natl Acad Sci U S A 109, 11717–11722.

Hagen, G., Müller, S., Beato, M., and Suske, G.J.T.E.j. (1994). Sp1-mediated transcriptional activation is repressed by Sp3. 13, 3843–3851.

Hirukawa, A., Smith, H.W., Zuo, D., Dufour, C.R., Savage, P., Bertos, N., Johnson, R.M., Bui, T., Bourque, G., and Basik, M.J.N.c. (2018). Targeting EZH2 reactivates a breast cancer subtype-specific antimetastatic transcriptional program. 9, 1–15.

Holm, K., Grabau, D., Lövgren, K., Aradottir, S., Gruvberger-Saal, S., Howlin, J., Saal, L.H., Ethier, S.P., Bendahl, P.-O., and Stål, O.J.M.o. (2012). Global H3K27 trimethylation and EZH2 abundance in breast tumor subtypes. 6, 494–506.

Hon, G.C., Hawkins, R.D., Caballero, O.L., Lo, C., Lister, R., Pelizzola, M., Valsesia, A., Ye, Z., Kuan, S., and Edsall, L.E.J.G.r. (2012). Global DNA hypomethylation coupled to repressive chromatin domain formation and gene silencing in breast cancer. 22, 246–258.

Hoy, S.M.J.D. (2020). Tazemetostat: First Approval. 1–9.

Hussein, Y.R., Sood, A.K., Bandyopadhyay, S., Albashiti, B., Semaan, A., Nahleh, Z., Roh, J., Han, H.D., Lopez-Berestein, G., and Ali-Fehmi, R.J.H.p. (2012). Clinical and biological relevance of enhancer of zeste homolog 2 in triple-negative breast cancer. 43, 1638–1644.

Innocente, S.A., and Lee, J.M.J.F.I. (2005). p53 is a NF-Y-and p21-independent, Sp1-dependent repressor of cyclin B1 transcription. 579, 1001–1007.

Jeng, K.S., Chang, C.F., and Lin, S.S. (2020). Sonic Hedgehog Signaling in Organogenesis, Tumors, and Tumor Microenvironments. Int J Mol Sci 21.

Joosse, S.A., Hannemann, J., Spotter, J., Bauche, A., Andreas, A., Muller, V., and Pantel, K. (2012). Changes in keratin expression during metastatic progression of breast cancer: impact on the detection of circulating tumor cells. Clin Cancer Res 18, 993–1003.

Karantza, V.J.O. (2011). Keratins in health and cancer: more than mere epithelial cell markers. 30, 127–138.

Karsli-Ceppioglu, S., Dagdemir, A., Judes, G., Lebert, A., Penault-Llorca, F., Bignon, Y.-J., and Bernard-Galion, D.J.S.R. (2017). The epigenetic landscape of promoter genome-wide analysis in breast cancer. 7, 1–8.

Kennecke, H., Yerushalmi, R., Woods, R., Cheang, M.C.U., Voduc, D., Speers, C.H., Nielsen, T.O., and Gelmon, K.J.J.o.c.o. (2010). Metastatic behavior of breast cancer subtypes. 28, 3271–3277.

Kim, J., Lee, Y., Lu, X., Song, B., Fong, K.-W., Cao, Q., Licht, J.D., Zhao, J.C., and Yu, J.J.C.r. (2018). Polycomb-and methylation-independent roles of EZH2 as a transcription activator. 25, 2808–2820.e2804.

Kim, K.H., and Roberts, C.W.J.N.m. (2016). Targeting EZH2 in cancer. 22, 128–134.

Kleer, C.G., Cao, Q., Varambally, S., Shen, R., Ota, I., Tomlins, S.A., Ghosh, D., Sewalt, R.G., Otte, A.P., and Hayes, D.F.J.P.o.t.N.A.o.S. (2003). EZH2 is a marker of aggressive breast cancer and promotes neoplastic transformation of breast epithelial cells. 100, 11606–11611.

Kolasinska-Zwierz, P., Down, T., Latorre, I, Liu, T., Liu, X.S., and Ahringer, J.J.N.g. (2009). Differential chromatin marking of introns and expressed exons by H3K36me3. 41, 376.

Lehmann, B.D., Bauer, J.A., Chen, X., Sanders, M.E., Chakravarthy, A.B., Shyr, Y., and Pietenpol, J.A.J.T.J.o.c.i. (2011). Identification of human triple-negative breast cancer subtypes and preclinical models for selection of targeted therapies. 121, 2750–2767.

Lui, V.W., Peyser, N.D., Ng, P.K., Hritz, J., Zeng, Y., Lu, Y., Li, H., Wang, L., Gilbert, B.R., General, I.J., et al. (2014). Frequent mutation of receptor protein tyrosine phosphatases provides a mechanism for STAT3 hyperactivation in head and neck cancer. Proc Natl Acad Sci U S A 111, 1114–1119.

Ma, A., Stratikopoulos, E., Park, K.-S., Wei, J., Martin, T.C., Yang, X., Schwarz, M., Leshchenko, V., Rialdi, A., and Dale, B.J.N.c.b. (2020). Discovery of a first-in-class EZH2 selective degrader. 16, 214–222.

Maheshwari, S., Avula, S.R., Singh, A., Singh, L.R., Palnati, G.R., Arya, R.K., Cheruvu, S.H., Shahi, S., Sharma, T., and Meena, S.J.M.C.T. (2017). Discovery of a Novel Small-Molecule Inhibitor that Targets PP2A-β-Catenin Signaling and Restricts Tumor Growth and Metastasis. 16, 1791–1805.

McDonald, O.G., Li, X., Saunders, T., Tryggvadottir, R., Mentch, S.J., Warmoes, M.O., Word, A.E., Carrer, A., Salz, T.H., and Natsume, S.J.N.g. (2017). Epigenomic reprogramming during pancreatic cancer progression links anabolic glucose metabolism to distant metastasis. 49, 367–376.

Nan, J., Wang, Y., Yang, J., and Stark, G.R. (2018). IRF9 and unphosphorylated STAT2 cooperate with NF-kappaB to drive IL6 expression. Proc Natl Acad Sci U S A 115, 3906–3911.

Oskarsson, T., Acharyya, S., Zhang, X.H., Vanharanta, S., Tavazoie, S.F., Morris, P.G., Downey, R.J., Manova-Todorova, K., Brogi, E., and Massague, J. (2011). Breast cancer cells produce tenascin C as a metastatic niche component to colonize the lungs. Nat Med 17, 867–874.

Papafotiou, G., Paraskevopoulou, V., Vasilaki, E., Kanaki, Z., Paschalidis, N., and Klinakis, A.J.N.c. (2016). KRT14 marks a subpopulation of bladder basal cells with pivotal role in regeneration and tumorigenesis. 7, 1–11.

Pekowska, A., Benoukraf, T., Ferrier, P., and Spicuglia, S.J.G.r. (2010). A unique H3K4me2 profile marks tissue-specific gene regulation. 20, 1493–1502.

Rothbart, S.B., and Baylin, S.B.J.C. (2020). Epigenetic Therapy for Epithelioid Sarcoma. 181, 211.

Schwartz, Y.B., and Pirrotta, V.J.N.R.G. (2007). Polycomb silencing mechanisms and the management of genomic programmes. 8, 9–22.

Seachrist, D.D., Sizemore, S.T., Johnson, E., Abdul-Karim, F.W., Weber Bonk, K.L., and Keri, R.A. (2017). Follistatin is a metastasis suppressor in a mouse model of HER2-positive breast cancer. Breast Cancer Res 19, 66.

Singh, A.K., Verma, A., Singh, A., Arya, R.K., Maheshwari, S., Chaturvedi, P., Nengroo, M.A., Saini, K.K., Vishwakarma, A.L., Singh, K., et al. (2020). Salinomycin inhibits epigenetic modulator EZH2 to enhance death receptors in colon cancer stem cells. Epigenetics, 1–18.

Varambally, S., Dhanasekaran, S.M., Zhou, M., Barrette, T.R., Kumar-Sinha, C., Sanda, M.G., Ghosh, D., Pienta, K.J., Sewalt, R.G., and Otte, A.P.J.N. (2002). The polycomb group protein EZH2 is involved in progression of prostate cancer. 419, 624–629.

Verma, A., Singh, A., Singh, A., Chaturvedi, P., Nengroo, M., and Datta, D. (2018). Epigenetic modulator EZH2 governs CSC properties and alters metastatic cascade. Paper presented at: CANCER MEDICINE (WILEY 111 RIVER ST, HOBOKEN 07030-5774, NJ USA).

Verma, A., Singh, A., Singh, A.K., Chaturvedi, P., Nengroo, M.A., kumar Saini, K., and Datta, D. (2020). Selective ezh2 functional activation alters the metastatic landscape of triple negative breast cancer (AACR).

Vishnoi, M., Liu, N.H., Yin, W., Boral, D., Scamardo, A., Hong, D., and Marchetti, D.J.M.o. (2019). The identification of a TNBC liver metastasis gene signature by sequential CTC-xenograft modeling. 13, 1913–1926.

Volk-Draper, L.D., Rajput, S., Hall, K.L., Wilber, A., and Rana, S.J.N. (2012). Novel model for basaloid triple-negative breast cancer: behavior in vivo and response to therapy. 14, 926–IN913.

Wolff, A.C., Hammond, M.E.H., Hicks, D.G., Dowsett, M., McShane, L.M., Allison, K.H., Allred, D.C., Bartlett, J.M., Bilous, M., Fitzgibbons, P.J.A.o.P., et al. (2014). Recommendations for human epidermal growth factor receptor 2 testing in breast cancer: American Society of Clinical Oncology/College of American Pathologists clinical practice guideline update. 138, 241–256.

Yin, X., Wolford, C.C., Chang, Y.S., McConoughey, S.J., Ramsey, S.A., Aderem, A., and Hai, T. (2010). ATF3, an adaptive-response gene, enhances TGF{beta} signaling and cancer-initiating cell features in breast cancer cells. J Cell Sci 123, 3558–3565.

Yomtoubian, S., Lee, S.B., Verma, A., Izzo, F., Markowitz, G., Choi, H., Cerchietti, L., Vahdat, L., Brown, K.A., and Andreopoulou, E.J.C.R. (2020). Inhibition of EZH2 Catalytic Activity Selectively Targets a Metastatic Subpopulation in Triple-Negative Breast Cancer. 30, 755–770. e756.

Yong, K.M.A., Ulintz, P.J., Caceres, S., Cheng, X., Bao, L., Wu, Z., Jiagge, E.M., and Merajver, S.D.J.S.r. (2020). Heterogeneity at the invasion front of triple negative breast cancer cells. 10, 1–9.

Young, M.D., Willson, T.A., Wakefield, M.J., Trounson, E., Hilton, D.J., Blewitt, M.E., Oshlack, A., and Majewski, I.J.J.N.a.r. (2011). ChIP-seq analysis reveals distinct H3K27me3 profiles that correlate with transcriptional activity. 39, 7415–7427.

