## Supplementary Figure for "H3K27me3 mediated KRT14 upregulation promotes TNBC peritoneal metastasis"

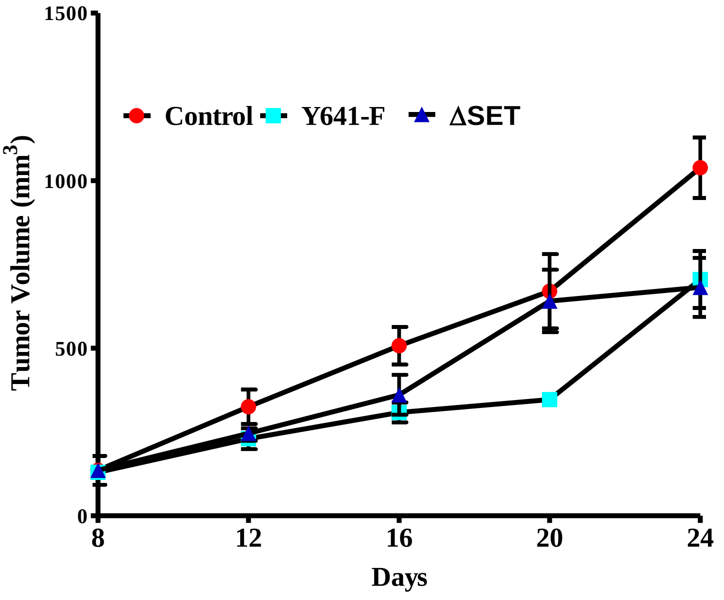


**Supplementary Figure :1**

**EZH2 (Y641-F ) and ΔSET OE have no significant impact on tumor growth.**

The control, EZH2 (Y641-F) OE 4T-1 and Δ SET ($1X10^6$) cells in 100 µl PBS were subcutaneous inoculated in the left flank of 4- to 6-week-old female nude Crl: CD1-Foxn1nude mice (n=5 each group) and allowed to grow for 25 days (n=5). The growth curve is shown; points are indicative of an average of tumor volume (error bar, +/- SEM).

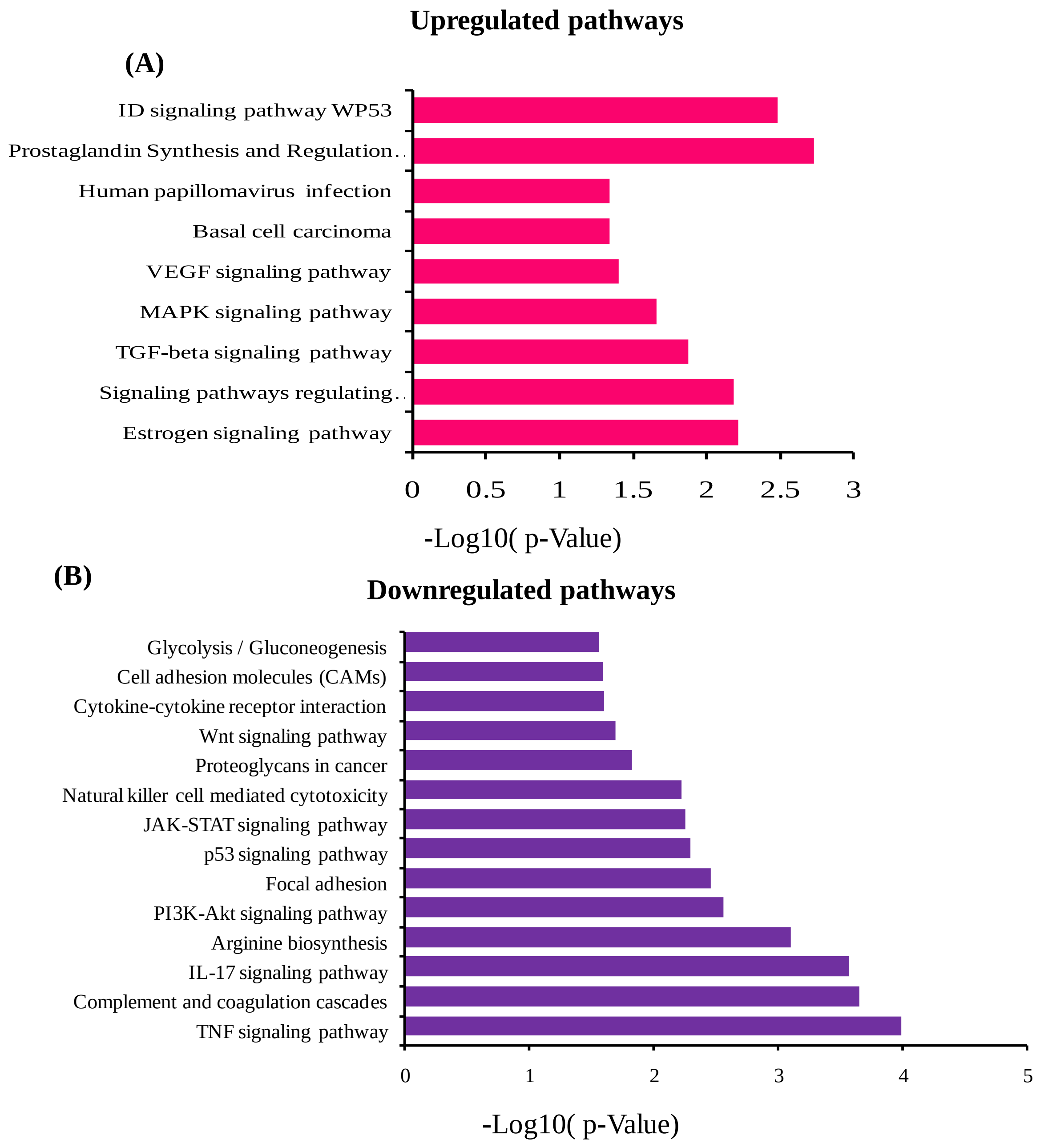


**Supplementary Figure: 2**

**KEGG pathway analysis of candidate genes:**

**(A and B)** RNA sequencing analysis was performed in Control and EZH2 Y641-F 4T1 cells. Details are described in method section. Significantly enriched KEGG pathways of unregulated and downregulated DEGs identified in EZH2(Y641-F) cells. ($p<0.05$) was considered to indicate a statistically significantly difference.

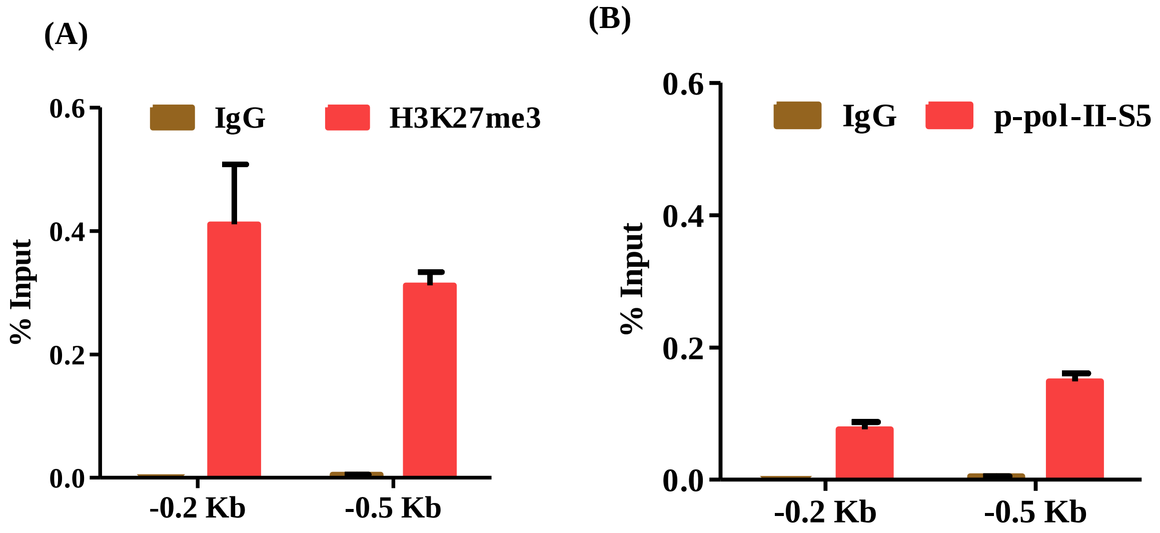


**Supplementary Figure: 3**

**The H3K27me3 and p-Pol-II-S5 enrichment analysis by ChIP-q PCR at *DLC1* promoter**

ChIP was performed in 4T-1 mouse cells using anti-H3K27me3, p-Pol-II-S5 and IgG antibodies and then examined by real-time q-PCR. using respective primers of *DLC1* gene. ChIP q-PRC results showing differential fold change in H3K27me3 and p-Pol-II-S5 enrichment at the promoter of *DLC1* gene. **(A and B)** The analysis of enrichment for H3K27me3 and p-Pol-II-S5 in the *DLC-1* Promoter at -0.2 Kb and -0.5 Kb regions from TSS respectively.

(A)

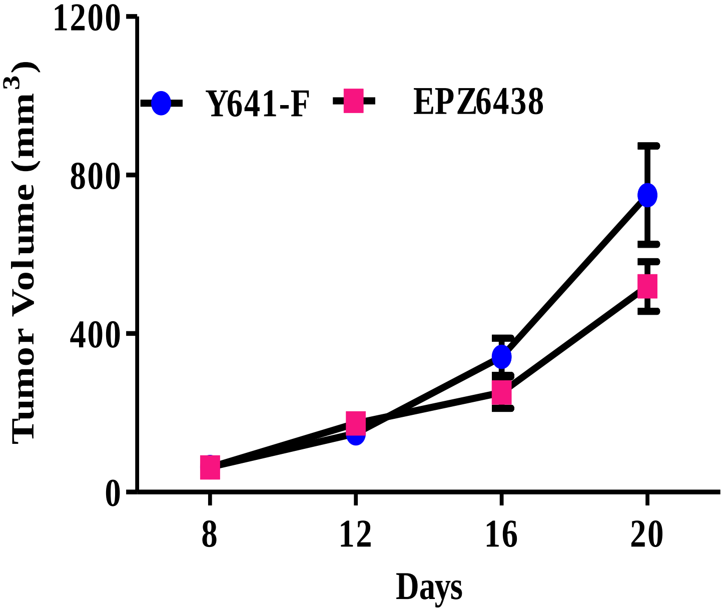


**Supplementary figure: 4**

**EPZ6438 treatment has no impact primary on TNBC tumor growth**

**(A)** 4T1 (Y641-F) cells ($1X10^6$) in 100 µl PBS were orthotopically inoculated in the left mammary fat pad of 4 to 6-week - old female nude and allowed to grow for 25 days. After 1 week mice were randomised and divided into vehicle and treatment groups (n=5). The EPZ6438 (250mg/Kg) dose was administered to mice once in a day by oral route. The growth curve analysis of control and EPZ6438 treated mice are shown, data points are indicative of an average of tumor volume (error bar, +/- SEM).

**Table S1. Upregulated biological KEGG pathways in 4T-1 EZH2 (Y641-F) cells predicated by Enricher Analysis**

| Term | P-value | Odds Ratio | Combined Score | Genes |
|---|---|---|---|---|
| Estrogen signaling pathway | 0.006034 | 5.527915976 | 28.24918404 | NCOA2;KRT16;KRT14;CREB5 |
| Signaling pathways regulating pluripotency of stem cells | 0.006519 | 5.406866721 | 27.21267894 | FZD5;ID2;ID3;SMAD5 |
| TGF-beta signaling pathway | 0.013192 | 6.105006105 | 26.42312327 | ID2;ID3;SMAD5 |
| MAPK Signaling Pathways | 0.0217 | 3.149407911 | 12.06364822 | HSPB1;FLNC;DUSP8;AREG;DUSP7 |
| VEGF signaling pathway | 0.039258 | 6.385696041 | 20.67437497 | HSPB1;PTGS2 |
| Basal cell carcinoma | 0.045588 | 5.878894768 | 18.15463266 | FZD5;PTCH1 |
| Human papillomavirus infection | 0.045684 | 2.572016461 | 7.937249223 | FZD5;MAML1;TNC;PTGS2;CREB5 |
| Prostaglandin Synthesis and Regulation WP98 | 0.001843 | 12.34567901 | 77.73572317 | ANXA1;SOX9;PTGS2 |
| ID signaling pathway WP53 | 0.0033 | 23.14814815 | 132.2666075 | ID2;ID3 |

**Table S2. Down regulated biological KEGG pathways in 4T-1 EZH2 (Y641-F) cells predicated by Enricher Analysis:**

| Term | P-value | Odds Ratio | Combined Score | Genes |
|---|---|---|---|---|
| TNF signaling pathway | 1.03E-04 | 6.526807 | 59.94707 | VCAM1;CSF1;CCL20;MMP3;CCL2;MMP9;ICAM1 |
| Complement and coagulation cascades | 2.22E-04 | 6.993007 | 58.82991 | C3;C1RA;C1S1;PROS1;ITGB2;PLAT |
| PI3K-Akt signaling pathway | 0.002728 | 2.872944 | 16.96192 | CSF3;CSF1;CCND1;COL6A1;PDGFB;SPP1;SGK1;PCK2;VEGFA;THBS3 |
| Focal adhesion | 0.003445 | 3.607783 | 20.45896 | CCND1;COL6A1;PDGFB;SPP1;PPP1R12B;VEGFA;THBS3 |
| p53 signaling pathway | 0.005119 | 5.778259 | 30.4795 | CCND1;GADD45A;SESN2;BID |
| JAK-STAT signaling pathway | 0.005498 | 3.752345 | 19.52475 | CSF3;CCND1;IFNGR2;PIM1;PDGFB;IRF9 |
| Natural killer cell mediated cytotoxicity | 0.006025 | 4.345937 | 22.21539 | IFNGR2;H2-K1;ITGB2;BID;ICAM1 |
| Proteoglycans in cancer | 0.014788 | 3.031451 | 12.77427 | WNT10A;CCND1;ANK2;PPP1R12B;MMP9;VEGFA |
| Wnt signaling pathway | 0.020428 | 3.205128 | 12.47072 | SFRP1;WNT10A;CCND1;CCN4;GPC4 |
| Cytokine-cytokine receptor interaction | 0.024741 | 2.458728 | 9.095561 | CSF3;CSF1;TNFSF15;CCL20;IFNGR2;TNFRSF9;CCL2 |
| Cell adhesion molecules (CAMs) | 0.025715 | 3.016591 | 11.04281 | ALDH3A1;ENO2;PCK2 |
| Pathways in cancer | 0.037421 | 1.917086 | 6.298638 | WNT10A;CCND1;NOS2;GADD45A;IFNGR2;PDGFB;PIM1;BID;MMP9;VEGFA |
| Glycolysis / Gluconeogenesis | 0.02772 | 4.592423 | 16.46651 | ALDH3A1;ENO2;PCK2 |
| Arginine biosynthesis | 7.88E-04 | 16.19433 | 115.721 | NOS2;GPT2;ASS1 |

**Table S3. List of Primers**

| Quantitative PCR (qPCR) Primers | | | |
|---|---|---|---|
| **S. No.** | **Gene Name** | **Primer Name (Mouse)** | **Sequence (5'-3')** |
| 1 | *FST* | qFST_FP | CATGGACCGATGGAGGATGT |
| | | qFST_RP | GCGGTAGGTTTTCCCATCCA |
| *2* | *DOK7* | qDOK7 _FP | CCAGCCACTCTTCACCTCTG |
| | | qDOK7_RP | AGCTCATCTGCTCTCCCTCA |
| 3 | *TLe6* | qTle6_FP | AGCAGCGTCGTCTCTGAAAT |
| | | qTle6_RP | CGGGGTGCTTTTGAACTTGG |
| 4 | *PTPRUC* | qPTRUC_FP | GCTGATGAGGGCAGACTGTT |
| | | qPTRUC_RP | TGCTCCCAGATCATCCTCCA |
| 5 | *DLC-1* | qDLC-1 _FP | GAAAGGCCACCACGAGAAGA |
| | | qDLC-1_RP | TCCTTTTCCGTACCATGGGC |
| 6 | *ATF-3* | qATF_3 FP | AAATTGCTGCTGCCAAGTGT |
| | | qATF-3 RP | TTCCGGTGTCCGTCCATTCT |
| 7 | *ID-2* | qID2_FP | CCGGTGAGGTCCGTTAGG |
| | | qID2_RP | CTGGACGCCTGGTTCTGTC |
| 8 | *FZD-5* | qFZD_FP | GGTGGCACTAAGACGGACAA |
| | | qFZD_RP | ATCCAGACTCCCGACGTGAT |
| 9 | *KRT14* | qKRT14_FP | CAGTCCCAGCTCAGCATGAA |
| | | qKRT14_RP | TGGGAAGATGAAAGGTGGGC |
| 10 | *ID3* | q ID3_FP | GCCTCTTAGCCTCTTGGACG |
| | | qID3_RP | CTGTCTGGATCGGGAGATGC |
| 11 | *DEPTOR* | qDEPTOR_FP | CGCATGACAGTCCCTTCTGT |
| | | qDEPTOR_RP | GCACAGATTTGGGGTTGCAG |
| 12 | *KRT16* | qKRT16_FP | TGCGCAAGGTGCTAGATGA |
| | | qKRT16_RP | GACCCCTCAAGGCAAGCATC |
| 13 | *KRT8* | qKRT8_FP | TGGAACTAGAGTCCCGCCTG |
| | | qKRT8_RP | TGCAACTCACGGATCTCCTC |

| S. No. | Gene Name | Primer Name (Human) | Sequence (5'-3') |
|---|---|---|---|
| 14 | *KRT14* | qKRT14_FP | TCTGAACGAGATGCGTGACC |
| | | qKRT14_RP | TGAAGAACCATTCCTCGGCA |

**Table S4: List of ChIP primers**

| Chromatin Immunoprecipitation-Quantitative PCR (ChIP-qPCR) Primers | | | |
|---|---|---|---|
| **S. No.** | **Gene Name** | **Primer Name** | **Sequence (5'-3')** |
| 1 | *KRT14 (mouse)* | Us_0.2Kb_FP | GGACGAGAAAGCCCAAAACAC |
| | | Us_0.2Kb_RP | CCCGATCAGATCCCTCCTCT |
| 2 | *KRT14 (mouse)* | Us_0.5 Kb_FP | CCAGCTAAGTGCCAGTCTCC |
| | | Us_0.5Kb_RP | AGTAGGGCCTTACCACACCA |
| 3 | *KRT14 (mouse)* | Us_1Kb_FP | GTTCCTCCTCCCCATACGTG |
| | | Us_1Kb_RP | TAAGGGCACATGCCTGGAAC |
| 4 | *KRT14 (mouse)* | Us_1.5Kb_FP | GGAAGGGTCAGGTGGGATTG |
| | | Us_1.5Kb_RP | GAGGCTCCTGCACTGTTCTT |
| 5 | *KRT14(Human)* | Us_0.2Kb_FP | CGGGACAAGAAAGCCCAAAA |
| | | Us_0.2Kb_RP | TATACTCGTGGGTAGGGGGC |
| 6 | *KRT14(Human)* | Us_0.7Kb_FP | GGGTGGGAACCACGATACAC |
| | | Us_0.7Kb_RP | ATGGATACCCGGCTGGAAAG |
| 7 | *KRT14(Human)* | Us_1.1Kb_FP | CAGTTCCACAAGGGGCTCAA |
| | | Us_1.1Kb_RP | AGAAGCCTCGTTGGCATTGT |
| 8 | *KRT14(Human)* | Us_1.5Kb_FP | TTTGCTGGCAGATTTGGGGA |
| | | Us_1.5Kb_RP | GCCTGACGCATCCTATCTCC |

**Table S5. List of ShRNA oligoes.**

| ShRNA Oligoes | | | |
|---|---|---|---|
| **S. No.** | **Gene Name** | **Primer Name** | **Sequence (5'-3')** |
| 1 | *KRT14 (Human)* | KRT14 _Sh1_FP | CCGGGCCTGCTGAGATCAAAGACTACTCGAGTAGTCTTTGATCTCAGCAGGCTTTTTG |
| | *KRT14 (Human)* | KRT14_Sh1_RP | AATTCAAAAAGCCTGCTGAGATCAAAGACTACTCGAGTAGTCTTTGATCTCAGCAGGC |
| 2 | *KRT14 (Human)* | KRT14_sh2_FP | CCGGGGTGCAGAGCGGCAAGAGCGACTCGAGTCGCTCTTGCCGCTCTGCACCTTTTTG |
| | | KRT14_sh2_RP | AATTCAAAAAGGTGCAGAGCGGCAAGAGCGACTCGAGTCGCTCTTGCCGCTCTGCACC |
| 3 | *KRT14 (Mouse)* | KRT14_Sh1_FP | CCGGCCAATTCTCCTCATCCTCTCACTCGAGTGAGAGGATGAGGAGAATTGG TTTTTG |
| | | KRT14_Sh1_RP | AATTCAAAAACCAATTCTCCTCATCCTCTCACTCGAGTGAGAGGATGAGGAGAATTGG |
| 4 | *KRT14 (Mouse)* | KRT14_Sh2_FP | CCGGGCCCACTGAGATCAAAGACTACTCGAG TAGTCTTTGATCTCAGTGGGC TTTTTG |
| | | KRT14_Sh2_RP | AATTCAAAAAGCCCACTGAGATCAAAGACTACTCGAGTAGTCTTTGATCTCAGTGGGC |
| 5 | *EZH2 (Mouse)* | EZH2_Sh1_FP | CCGGGCACAAGTCATCCCGTTAAAGCTCGAGCTTTAACGGGATGACTTGTGCTTTTTG |
| | | EZH2_Sh1_RP | AATTCAAAAACACAAGTCATCCCGTTAAAGCTCGAGCTTTAACGGGATGACTTGTGC |
